# STK19 facilitates the clearance of lesion-stalled RNAPII during transcription-coupled DNA repair

**DOI:** 10.1101/2024.07.22.604575

**Authors:** Diana van den Heuvel, Marta Rodríguez-Martínez, Paula J. van der Meer, Nicolas Nieto Moreno, Jiyoung Park, Hyun-Suk Kim, Janne J.M. van Schie, Annelotte P. Wondergem, Areetha D’Souza, George Yakoub, Anna E. Herlihy, Krushanka Kashyap, Thierry Boissière, Jane Walker, Richard Mitter, Katja Apelt, Klaas de Lint, Idil Kirdök, Mats Ljungman, Rob M.F. Wolthuis, Patrick Cramer, Orlando D. Schärer, Goran Kokic, Jesper Q Svejstrup, Martijn S. Luijsterburg

## Abstract

Transcription-coupled DNA repair (TCR) removes bulky DNA lesions impeding RNA polymerase II (RNAPII) transcription. Recent studies have outlined the stepwise assembly of TCR factors CSB, CSA, UVSSA, and TFIIH around lesion-stalled RNAPII. However, the mechanism and factors required for the transition to downstream repair steps, including RNAPII removal to provide repair proteins access to the DNA lesion, remain unclear. Here, we identify STK19 as a new TCR factor facilitating this transition. Loss of STK19 does not impact initial TCR complex assembly or RNAPII ubiquitylation but delays lesion-stalled RNAPII clearance, thereby interfering with the downstream repair reaction. Cryo-EM and mutational analysis reveal that STK19 associates with the TCR complex, positioning itself between RNAPII, UVSSA, and CSA. The structural insights and molecular modeling suggest that STK19 positions the ATPase subunits of TFIIH onto DNA in front of RNAPII. Together, these findings provide new insights into the factors and mechanisms required for TCR.

## Introduction

Elongating RNA polymerase II (RNAPII) acts as the damage sensor to initiate transcription-coupled DNA repair (TCR), a sub-pathway of nucleotide excision repair (NER) that resolves helix-distorting DNA damage from the template DNA strand in genes (Bohr et al., 1985; Mellon et al., 1987). The stalling of RNAPII triggers the initial recruitment of TCR protein (Xu et al., 2017). However, stalled RNAPII shields the DNA lesion and prevents downstream NER proteins from accessing and removing the DNA damage. Consequently, the processing and removal of RNAPII from the lesion must be closely linked with the progression of TCR. How the RNAPII-bound complex that shields the DNA lesion transitions to downstream repair steps by NER proteins remains unresolved (Jia et al., 2021; Lans et al., 2019; Nieto Moreno et al., 2023; van den Heuvel et al., 2021b; Wang et al., 2023).

TCR is initiated by the sequential and cooperative association of the TCR-specific proteins CSB, CSA, and UVSSA with DNA damage-stalled RNAPII bound to elongation factor ELOF1 that further positions TCR proteins on the surface of RNAPII (Geijer et al., 2021; Kokic et al., 2024; van der Weegen et al., 2021; van der Weegen et al., 2020). Ultimately, the TCR factors mediate the recruitment of transcription factor IIH (TFIIH) to RNAPII, a general repair factor shared between TCR and the global genome repair (GGR) sub-pathway of NER. The TFIIH core complex consists of a seven-subunit core composed of XPB, XPD, p62, p52, p44, p34 and p8/TTDA (Compe and Egly, 2016). XPB is an ATP-dependent 3’ to 5’ translocase that binds DNA downstream of RNAPII during transcription initiation and facilitates promoter opening. In contrast, XPD is an ATP-dependent 5’ to 3’ helicase that is dispensable during transcription initiation and is not engaged with DNA during this process (He et al., 2016). Rearrangement of the TFIIH structure activates the complex for repair by enabling both ATPase subunits to bind DNA (Kokic et al., 2019). Upon such activation, TFIIH triggers the formation of an open DNA bubble and verifies the presence of a bulky DNA lesion needed for the assembly of the NER preincision complex (Coin et al., 2008; Fu et al., 2023; Fu et al., 2022; Kim et al., 2023; Kokic et al., 2019). Downstream NER proteins, including XPA, RPA, and the endonucleases XPG and ERCC1-XPF, subsequently mediate dual incision around the lesion, followed by removal of the damaged DNA strand and DNA gap-fill synthesis of the single-stranded gap followed by ligation (Aboussekhra et al., 1995).

During GGR, the XPC complex recruits TFIIH to the lesion, where only XPB initially binds DNA, while the XPD repair helicase is located away from DNA (Kim et al., 2023), similar to transcription initiation (Aibara et al., 2021; He et al., 2016). Upon binding, XPA wedges between the XPB and XPD subunits and clamps the DNA, reorienting the complex such that XPD also engages DNA (Kim et al., 2023; Kokic et al., 2019). Thus the association of downstream NER proteins activates TFIIH for repair in GGR (Coin et al., 2008; Kokic et al., 2019). However, during TCR, the large lesion-stalled RNAPII prevents the recruitment of downstream NER proteins to the TCR complex already containing TFIIH (van der Weegen et al., 2020). This suggests that there must be a critical transition from TFIIH bound to RNAPII decorated with TCR factors (possibly in a transcription initiation-like state) to TFIIH with both ATPase subunits engaged with DNA (in the repair state) that would enable the recruitment of downstream NER proteins. This transition must involve the removal of RNAPII, thereby providing access to the DNA lesion.

The importance of timely removal of stalled RNAPII during TCR is underscored by clinical manifestations in Cockayne Syndrome (CS) patients, who exhibit persistent RNAPII stalling at DNA lesions, resulting in severe neurological dysfunction and a shortened lifespan (Nance and Berry, 1992; van den Heuvel et al., 2021b). Two possibly intertwined mechanisms have been suggested to regulate RNAPII removal. First, the early TCR factors, including CSB, CSA, and ELOF1, promote the ubiquitylation of a single lysine on the largest subunit of RNAPII (RPB1-K1268) (Nakazawa et al., 2020; Tufegdzic Vidakovic et al., 2020). Displacement of RNAPII from the site of DNA damage through TCR-mediated ubiquitylation may be part of the ‘last resort’ pathway that degrades stalled RNAPII when other options have failed (Wilson et al., 2013). Alternatively, or additionally, the activated TFIIH complex may remodel RNAPII and trigger its displacement from DNA lesions in an ATP-dependent manner (Sarker et al., 2005). This model would require correct placement and activation of TFIIH around lesion-stalled RNAPII prior to the recuritment of core NER proteins. However, which factors regulate RNAPII clearance during TCR is not known. Here, we identify the STK19 protein as an integral part of the TCR complex required for the timely removal of RNAPII from DNA lesions to allow downstream DNA repair.

## Results

### Multiple genetic screens identify STK19 as a putative TCR factor

Previous genome-wide screens led to the identification of numerous proteins involved in the cellular response to bulky DNA lesions (Boeing et al., 2016; Geijer et al., 2021; Olivieri et al., 2020; van der Weegen et al., 2021). To specifically identify new TCR genes, we previously performed a genome-wide CRISPR screen in the presence of Illudin S (van der Weegen et al., 2021), but a shortcoming of this approach is that Illudin S generates a variety of DNA lesions, many of which are not repaired by TCR. Indeed, while validating screen hits, we observed that several gene knockouts, such as *HMCES* and *XRCC1*, conferred sensitivity to Illudin S without affecting transcription recovery after UV irradiation, which is a hallmark of TCR (Mayne and Lehmann, 1982) (Figure S1a-f). In addition, knockout of *HMCES* or *XRCC1* in a *CSB*-deficient background caused additive Illudin S sensitivity (Figure S1b,e). Thus, Illudin S sensitization alone did not enable us to unequivocally classify new TCR genes.

To classify TCR genes with more confidence, we performed a second genome-wide CRISPR screen in the presence of the compound 4-nitroquinoline 1-oxide (4-NQO) (Figure S2a,b), which induces UV-like DNA lesions that are repaired by NER. Overlaying the two screens revealed genes that sensitize cells to either 4-NQO (*KEAP1*, *CUL3*) or Illudin S (*HMCES*, *XRCC1*) (Figure S2b). Importantly, we found that all genes that scored highly in both genetic screens encoded TCR factors, including *CSA*, *UVSSA* and *ELOF1*. We also analyzed the overlap between these genome-wide CRISPR screens and our previous siRNA screen for transcription recovery after UV (Figure S2c) (Boeing et al., 2016), as well as with relevant DNA damage response screens performed by (Olivieri et al., 2020). A recurring hit at the intersection between all these screens is the *STK19* gene (Figure S2d), which we therefore decided to study in detail.

### STK19-deficient cells are sensitive to transcription-blocking DNA damage

As part of our initial work on this protein, we previously showed that the *STK19* (Serine Threonine Kinase 19) gene has been incorrectly annotated in the human genome (Rodriguez-Martinez et al., 2020). Indeed, in spite of its name, STK19 is not a kinase but encodes a 254 amino acid chromatin-associated protein (Rodriguez-Martinez et al., 2020), with a poorly understood role in the cellular response to DNA damage (Boeing et al., 2016; Olivieri et al., 2020). To characterize STK19’s potential role in TCR, we generated knockout cells in four different genetic backgrounds, which were confirmed by sequencing and western blot analysis (Figure S3a-d). Consistent with the genetic screens, knockout of *STK19* caused strong sensitivity to different types of transcription-blocking DNA damage triggered by UV irradiation, Illudin S, 4-NQO, or cisplatin in all genetic backgrounds, although not to the same extent as cells with knockout of *CSB* or *ELOF1* (Figure 1a-d, Figure S3e-g). We note that while not all experiments were performed in all four genetic backgrounds, the results obtained were, without exception, consistent between the different cell lines tested, attesting to the generality of the conclusions made below.

**Figure 1.**
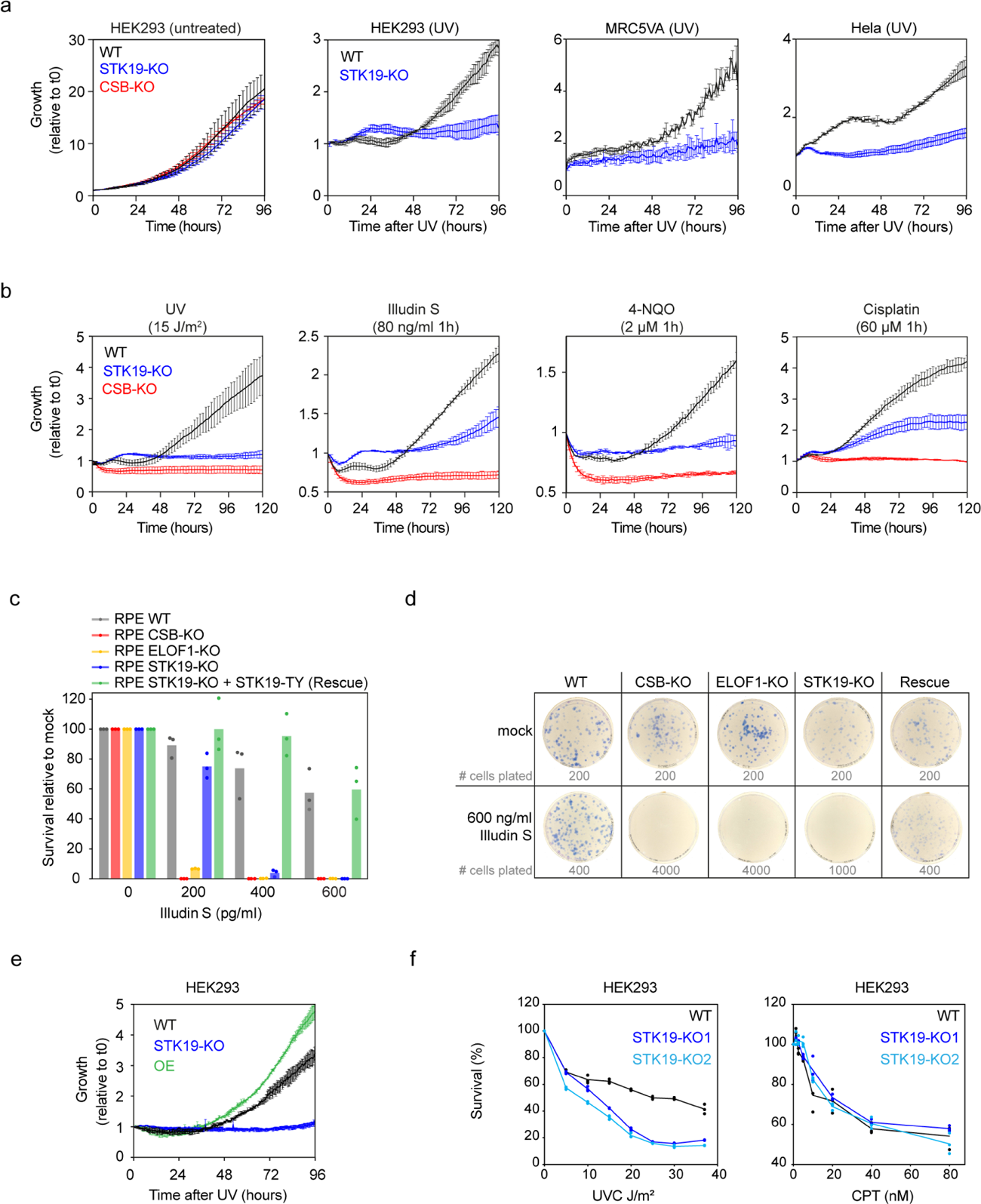
*STK19*-deficient cells are sensitive to transcription-blocking DNA damage. **a.** Quantification of cell growth using Incucyte, with or without 15 J/m^2^ UV-C irradiation in the wild type and mutant cell lines indicated. Cell growth (confluency) was monitored every 3 h and the data were normalized to t = 0 for each well. Data are represented at each 3 h time point as average relative confluency of 2 biological replicates ± SD. Experiments were repeated 3 times. **b.** As in A, but also after 1 hour treatment with the indicated compounds. **c.** Quantification of clonogenic survival after Illudin S treatment in RPE1 cells with the indicated genotype. Bars represent the means of 3 experiments with individual data points shown as circles. **d.** Representative examples of clonogenic growth in the presence of Illudin S as in **c**. **e.** As in A but also showing STK19 over-expressing cells (OE). Data are represented at each 3 h time point as average relative confluency of 2 biological replicates ± SD. Experiments were repeated 3 times with different UV doses. **f.** Quantification cell survival after irradiation with increasing doses of UV-C (left) or after 1 hour treatment with increasing doses of camptothecin (CPT, right) in the indicated cell lines. Cell growth was quantified using luminescence CellTiter-Glo assay. Lines represent the means of 3 experiments with individual data points shown as circles.

Importantly, only sgRNAs targeting the correctly annotated 254 amino acid protein resulted in sensitivity to Illudin S, while an sgRNA targeting the mis-annotated genomic region preceding the correct start codon of *STK19* did not (Figure S3h). Indeed, the sensitivity of *STK19-*KO cells to Illudin S could be rescued by re-expressing a cDNA encoding the 254 amino acid STK19 protein in RPE1 cells (Figure 1c,d). Moreover, over-expression of STK19 in HEK293 cells even provided some resistance to UV irradiation compared to WT cells (Figure 1e).

DNA lesions triggered by UV light, 4-NQO and cisplatin activate both the GGR and the TCR pathways of nucleotide excision repair. To measure a potential involvement of STK19 in GGR, we measured gap-fill DNA synthesis by 5-ethynyl-2-deoxyuridine (EdU) incorporation at sites of local UV-induced DNA damage (van der Meer et al., 2023). Both WT and *STK19*-deficient cells exhibited robust GGR-dependent DNA synthesis, while *XPC*-deficient cells, included for comparison, displayed a deficiency in this process, as expected (Figure S3i,j). This indicates that STK19 is not involved in GGR. Importantly, *STK19*-deficient cells are neither sensitive to replication stress nor DNA breaks as shown by exposure to camptothecin (CPT), hydroxyurea (HU), or neocarzinostatin (NCS) (Figure 1f, Figure S3k,l). Together, these data specifically implicate STK19 in the response to transcription-blocking DNA damage that is repaired by TCR.

### STK19 regulates transcription recovery

A hallmark of TCR deficiency is the inability to resume transcription after UV irradiation, resulting in bulky lesions that block the progression of RNAPII (Mayne and Lehmann, 1982). To measure transcription resumption after UV irradiation, we metabolically labelled and visualized nascent transcripts by 5-ethynyl-2-uridine (EU) incorporation followed by microscopy (Figure 2a,b). All cell lines showed a strong drop at 3 h after UV, which is the time point where we typically observe the maximum decrease in nascent transcription. At 16 h, *STK19-*KO cells failed to resume transcription to the levels observed in WT cells, in a manner that was rescued by re-expression of STK19 (Figure 2b). We note that compared to the *CSB*-deficient cells tested in parallel, low levels of residual transcription recovery were detected in the *STK19*-KO cell line, suggesting that TCR is not completely inactivated in *STK19*-deficient cells.

**Figure 2.**
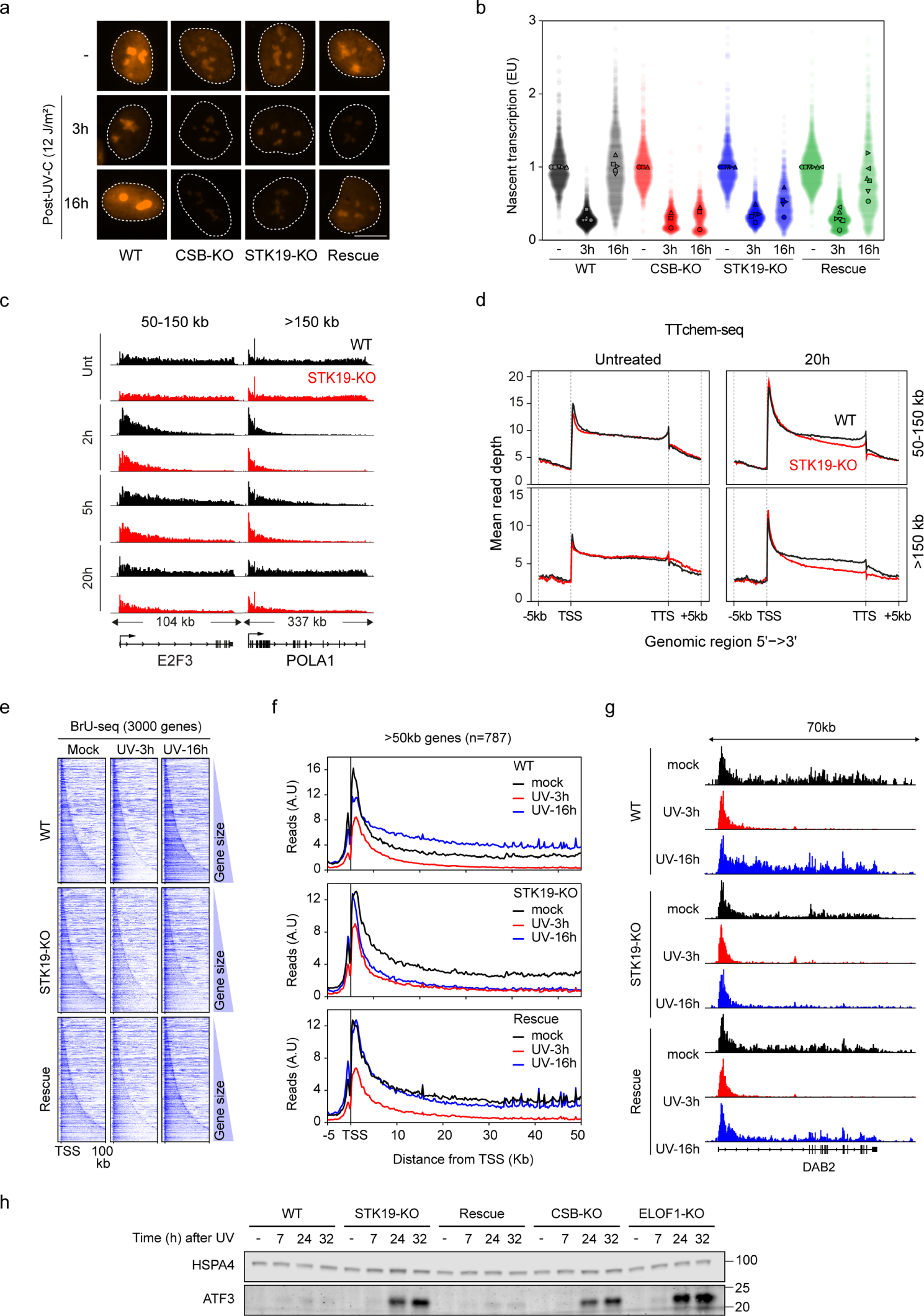
STK19 promotes transcription recovery in response to DNA damage. **a.** Microscopy images of the indicated RPE1 cells after 1 h EU-labelling in untreated (-) condition, or 3 h and 16 h after irradiation with 12 J/m^2^ UV-C. Scale bar, 10 μm. **b.** Quantification of **a**. All cells are depicted as individual data points with the bar representing the mean of all data points. The individual means of five biological replicates are depicted as open symbols. **c.** Browser tracks from TTchem-seq experiment in untreated cells and 2 h and 20 h after UV irradiation (15 J/m^2^). Represented are genes E2F3 and POLA1. The data are normalized to yeast spike-in. **d.** Metagene TTchem-seq profiles of genes 50-150 kb and ≧150 kb, in untreated cells and 20h after UV irradiation (15 J/m^2^). Data are normalized to yeast spike-in. TSS, transcription start site. TTS, transciption termination sites. **e.** Heatmaps of BrU-seq data in RPE1 cells with the indicated genotype from the TSS into the first 100 kb of 3000 genes with strongest BrU signal in mock-treated cells, sorted on gene length. Heatmaps of the same genes at 3 or 16 h after UV-C irradiation with 12 J/m^2^. **f.** Average read count profiles from -5 kb to +50 kb around the transcription start site (TSS) of 787 genes above 50 kb in RPE1 cells with the indicated genotype that were untreated or at 3 or 16 h after UV-C irradiation with 12 J/m^2^. **g.** BrU-seq density across the *DAB2* gene in RPE1 cells with the indicated genotype that were untreated or at 3 or 16 h after UV-C irradiation with 12 J/m^2^. **h.** Western blot analysis of ATF3 protein level in RPE1 cells with the indicated genotype that were untreated or at 7, 24 or 32 h after UV-C irradiation with 12 J/m^2^. HSPA4 serves as a loading control.

To extend these findings, we performed genome-wide sequencing of nascent transcripts using two complementary approaches in different genetic backgrounds. Sequencing of 4-thiouridine (4-SU)-labelled nascent transcripts from HEK293 cells using a TT_chem_-seq protocol (Gregersen et al., 2020) showed no apparent difference between untreated WT and *STK19*-deficient cells by analysis before and 2 h after UV (Figure 2c,d). In both, nascent transcription progressively decreased across gene bodies 2 h after UV, in line with the frequency of photolesions and the probability of RNAPII running into them (Andrade-Lima et al., 2015; Perdiz et al., 2000; Tufegdzic Vidakovic et al., 2020; Williamson et al., 2017). However, in contrast to WT cells, *STK19*-deficient cells showed clear defects in transcription restart after UV, which was most striking 20 h after UV towards the 3’-end of long genes (Figure 2c,d). We confirmed these results by RNA RT-qPCR for the long *POLA1* and *EXT1* genes (Figure S4a).

Similarly, capturing BrU-labelled nascent transcripts from RPE1 cells using a BrU-seq protocol (Paulsen et al., 2014) also revealed little or no difference between untreated cells and a strong decrease in nascent transcription further into gene bodies in both WT and *STK19*-deficient cells at 3 h after UV irradiation. At 16 h after UV, we detected full transcription recovery in WT cells shown in heatmaps for 3000 genes sorted by gene length, or by metagene-analysis for 787 genes of at least 50 kb in length (Figure 2e,f,g). By contrast, *STK19*-deficient cells showed a clear transcription recovery defect, which was most easily seen in genes >50kb (Figure 2f, Figure S4b). Re-expression of STK19 rescued the transcription recovery defect (Figure 2f). At late time-points after UV, we observed elevated ATF3 protein levels, a marker of persistent transcription stress (Epanchintsev et al., 2017; Kristensen et al., 2013), to levels also observed in other TCR-deficient cells, which was again rescued by re-expression of STK19 (Figure 2h). Together, these results show that STK19 is required for the resumption of transcription in response to transcription-blocking DNA lesions.

### STK19 is required for TCR-dependent gap-fill DNA synthesis

Deficient transcription recovery may result from defective DNA repair, or from impaired transcription restart after repair. We therefore investigated whether STK19 is necessary for the successful execution of TCR-dependent DNA repair. After UV irradiation, TCR-dependent unscheduled gap-fill DNA synthesis (TCR-UDS) makes only a minor contribution to the overall cellular NER activity, which mainly reflects GGR. To eliminate GGR-dependent UDS and specifically measure TCR-UDS, we locally UV irradiated serum-starved *XPC-*KO cells, followed by incubation with ethynyl-deoxy-uridine (EdU) (van den Heuvel et al., 2023) (Figure 3a,b). As expected, the UDS signal disappeared when *CSB* was also knocked out in these cells, showing that the EdU incorporation observed in *XPC*-KO cells is TCR specific (Figure 3a,b). We subsequently generated *XPC*/*STK19* double knockout cells (Figure S3b), and also re-expressed TY1-STK19 in two independent dKO clones as a control. A strong TCR-UDS defect was observed in *STK19*-deficient cells, which was rescued by re-expression of STK19 (Figure 3a,b). These findings indicate that STK19 indeed has an important role in the TCR pathway.

**Figure 3.**
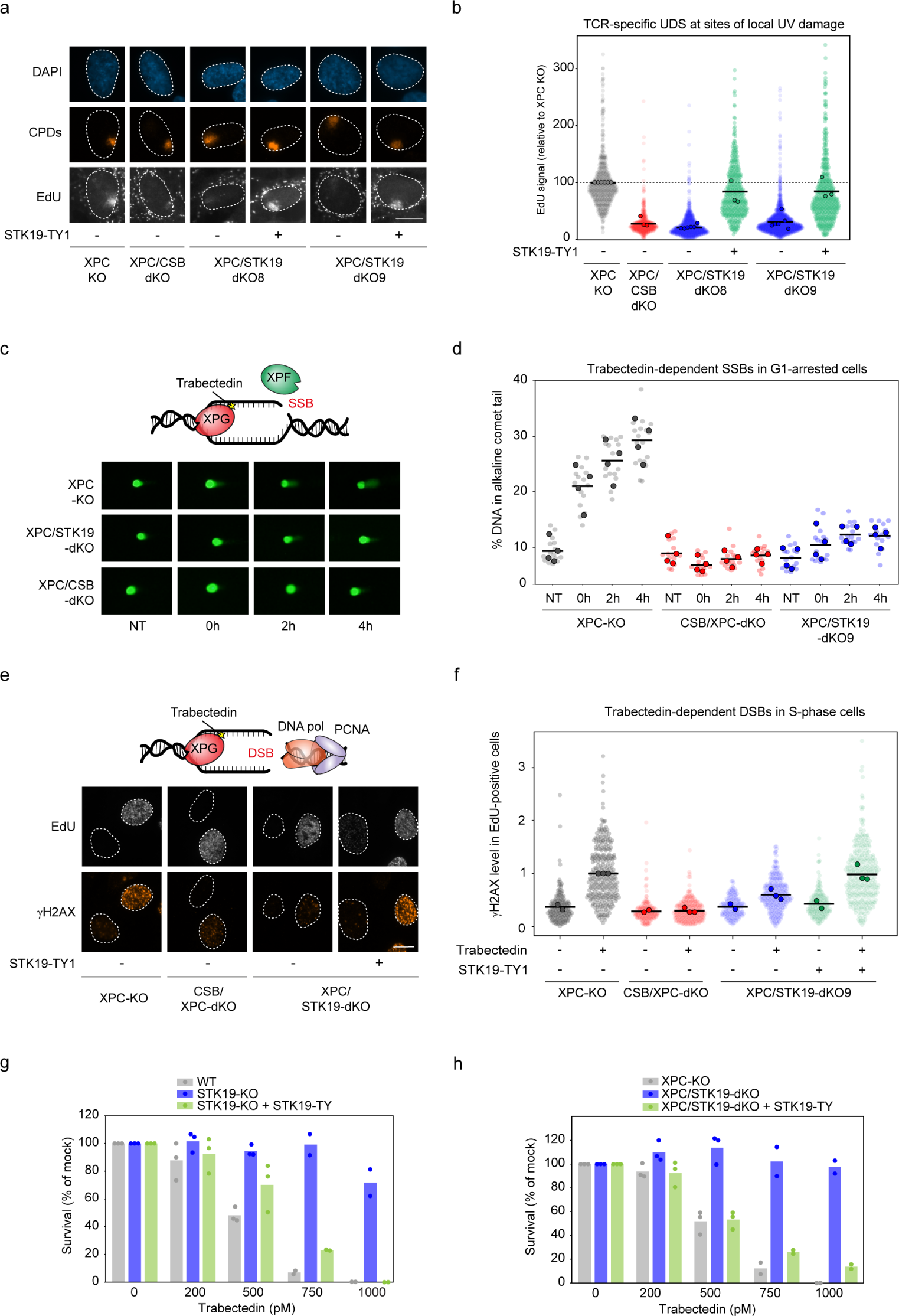
STK19 promotes downstream DNA repair steps. **a.** Representative images of EdU incorporation in RPE1 cells deficient in GGR (*XPC-*KO cells) with the indicated genotype to specifically measure UDS by TCR after 100 J/m^2^ UV-C irradiation through 5 µm pore membranes. Sites of local damage are identified by CPD staining. Scale bar, 10 μm. **b.** Quantification of TCR-UDS signal from **a**. All cells are depicted as individual semi-transparent data points with the bar representing the mean of all data points. The individual means of three to five biological replicates are depicted as solid circles with black lines. **c.** Representative images of 30 μm COMET chip from G1-arrested RPE1 cells deficient in GGR (*XPC-* KO cells) with the indicated genotype that were either untreated (NT) or treated for 2 h with 50 nM trabectedin and subsequently released for 0, 2, or 4 h. **d.** Quantification of **c**. The means of all technical replicates are depicted as individual semi-transparent data points with the bar representing the mean of all data points. The individual means of four biological replicates (with four technical replicates per experiment) are depicted as solid circles with black lines. **e.** Representative images of TCR-dependent γH2AX induction following trabectedin exposure in replicating cells labelled with 5-ethynyl-deoxyuridine (EdU) in RPE1 cells deficient in GGR (*XPC-*KO cells) with the indicated genotype. Scale bar, 10 μm. **f.** Quantification of **e**. The γH2AX levels were normalized to the average of the trabectedin-treated parental cells within each experiment. All cells are depicted as individual semi-transparent data points with the bar representing the mean of all data points. The individual means of three biological replicates are depicted as solid circles with black lines. **g, h.** Quantification of clonogenic survival after Trabectedin treatment in (**g**) RPE1 cells or (**h**) *XPC-*KO RPE1 cells with the indicated genotype. Bars represent the means of 3 experiments with individual data points shown as colored circles.

### STK19 is necessary for TCR-dependent incisions

The DNA synthesis step in TCR is preceded by a dual incision on each side of the lesion by the endonucleases ERCC1-XPF and XPG to excise a damaged, single-stranded DNA fragment (Marteijn et al., 2014). To further investigate where in the TCR mechanism STK19 acts, we used the chemotherapeutic agent trabectedin, which is specifically toxic to TCR-proficient cells (Takebayashi et al., 2001). Trabectedin-generated DNA lesions are recognized and processed by TCR proteins, resulting in an abortive repair intermediate consisting of a persistent single-strand break (SSBs), which is generated by an uncoupled ERCC1-XPF incision, with the DNA adduct blocking the subsequent XPG incision (Son et al., 2024). Such SSBs are detrimental to cell survival but are absent in TCR-deficient cells, which therefore uniquely survive trabectedin treatment.

Using a high-throughput alkaline COMET assay (Son et al., 2024), we detected an increased tail moment, reflecting TCR-dependent single incisions in response to trabectedin treatment in G1-synchronized RPE1 cells (Figure 3c,d). The trabectedin-induced SSBs increased 3-fold within the first 4 h after treatment in WT cells, while CSB-deficient cells generated few or no detectable SSBs in this assay. Similarly, STK19-deficient cells showed strongly suppressed SSB formation after trabectedin treatment (Figure 3c,d), strongly suggesting that the first ERCC1-XPF mediated incision of the TCR reaction does not take place in these cells either.

To further corroborate these results, we allowed cells with persistent SSBs to go through replication before measuring DNA double-strand break (DSB) formation by γH2AX staining, specifically in EdU-positive cells. To exclude interference from GGR, we again performed all experiments in an *XPC*-deficient background. *XPC-*KO (control) cells showed strongly elevated γH2AX in replicating cells after 4 h trabectedin treatment (Figure 3e,f), in line with the notion that TCR-dependent incisions cause SSBs that can subsequently be converted into DSBs during DNA replication. By contrast, CSB-deficient cells showed little or no such DNA breakage in replicating cells, and breakage was also strongly suppressed in *STK19*-deficient cells, in line with the results from the alkaline COMET assay. Re-expression of STK19 protein brought the trabectedin-induced γH2AX signal back to WT levels (Figure 3e,f). Clonogenic survival assays confirmed these findings and showed that both WT and *XPC*-deficient cells are sensitive to trabectedin in a dose-dependent manner. By contrast, *STK19*-KO rendered cells resistant to trabectedin in both WT and *XPC*-deficient genetic backgrounds, while re-expression of STK19 again sensitized cells to trabectedin (Figure 3g,h). Together, these results support the conclusion that STK19 is a new core TCR factor required for reaching the DNA incision step during TCR.

### STK19 associates with the RNAPII–bound TCR complex without affecting its assembly

Having established that STK19 is a new TCR factor, we asked where this protein acts during repair. TCR proteins associate with DNA damage-stalled RNAPII, culminating in TFIIH recruitment to somehow initiate the downstream nucleotide excision repair reaction (Kokic et al., 2021; Kokic et al., 2024; Nakazawa et al., 2020; van der Weegen et al., 2021; van der Weegen et al., 2020). To investigate whether STK19 has a role in TCR complex assembly, we immunoprecipitated RNAPII after UV irradiation to isolate TCR complexes (van der Weegen et al., 2020). Both CSB and CSA associated with RNAPII in a UV-dependent manner in all cell-lines tested (Figure 4a), showing that STK19 is not required for this early TCR step. The UV-induced association of TFIIH with RNAPII was slightly attenuated in *STK19*-deficient cells, while it was strongly reduced in the *ELOF1*-deficient cells tested in parallel (van der Weegen et al., 2021) (Figure 4a). The slightly decreased recruitment of TFIIH seems unlikely to, in itself, cause the strong DNA damage phenotypes observed for *STK19*-KO cells, but could suggest an incorrect positioning or decreased stability of TFIIH within the TCR complex. Re-expression of TY1-tagged STK19 rescued the attenuated TFIIH recruitment in *STK19*-KO cells. With the ability to detect the STK19 protein through the TY1 tag, we noticed that it co-immunoprecipitated with RNAPII in a UV-stimulated manner (Figure 4a). Independent RNAPII co-immunoprecipitation experiments in RPE1 cells confirmed this (Figure 4b), and we reciprocally found that RNAPII can co-immunoprecipitate with STK19 from HeLa cells (Figure S4c). We conclude that STK19 associates with the RNAPII-bound TCR complex without markedly affecting its initial assembly.

**Figure 4.**
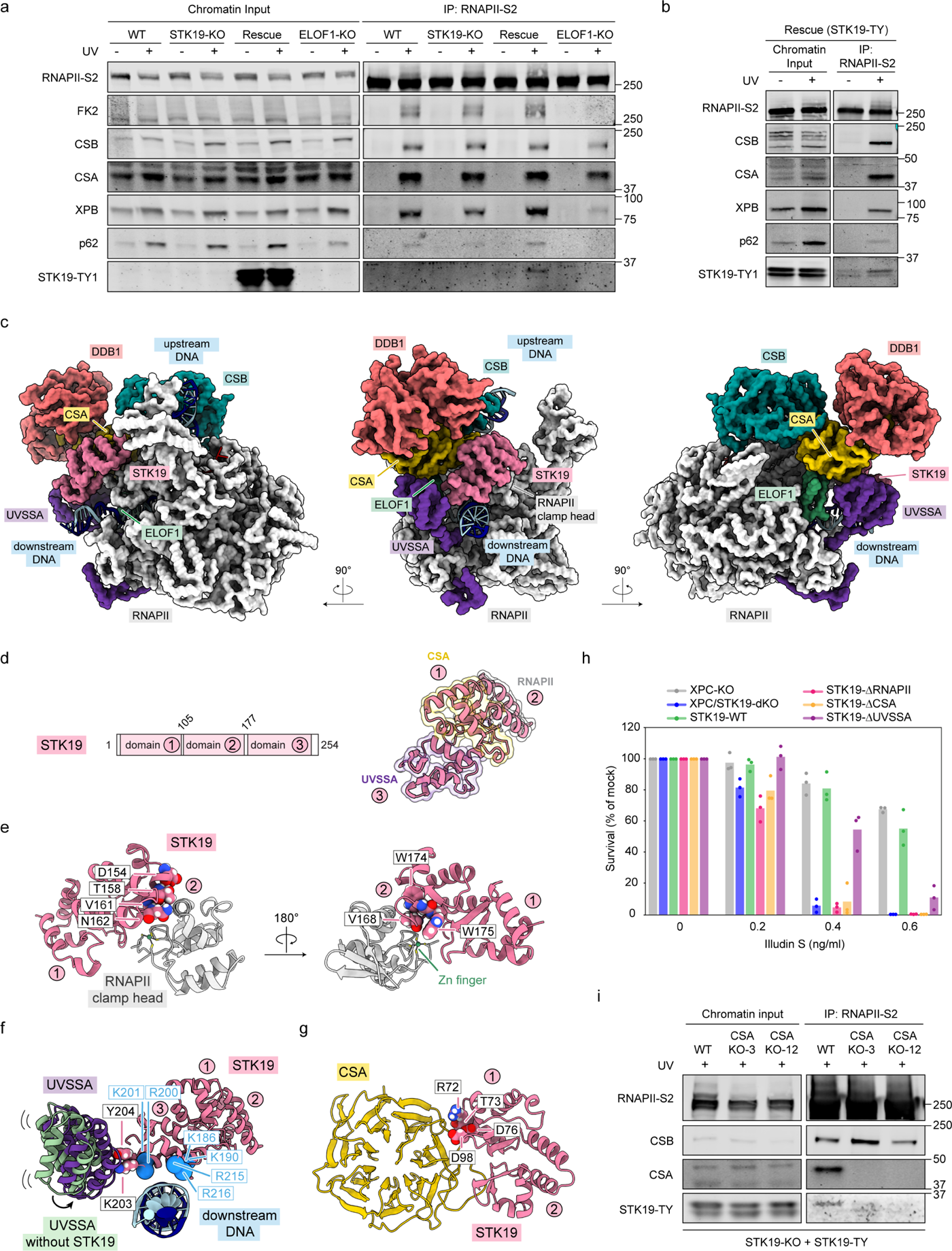
Structure of the RNAPII-TCR-STK19 complex. **a.** Endogenous RNAPII-S2 immunoprecipitation (IP) in RPE1 cells with the indicated genotype that were untreated or following UV irradiation (12 J/m^2^, 1 h recovery). TCR complex assembly and RNAPII ubiquitylation were analyzed with the indicated antibodies. The data shown represent at least three independent experiments. **b.** Independent Co-IP experiment as shown in **a**. **c.** Molecular model of the RNAPII-TCR-STK19 complex. **d.** Schematics of the STK19 domain composition **e.** Zoom-in on the STK19-RNAPII interface. STK19 residues at the interface are shown as spheres. **f.** Zoom-in on the STK19-UVSSA interface. UVSSA in the RNAPII-TCR complex without STK19 is shown in green and was modelled based on our previous structure (Kokic et al., 2024) by aligning structures on CSA. Positively charged STK19 residues in the vicinity of downstream DNA are shown as blue spheres. **g.** Zoom-in on the STK19-CSA interface. STK19 residues at the interface are shown as spheres. **h.** Quantification of clonogenic survival after Illudin S treatment in RPE1 cells with the indicated genotype. The following amino acid substitutions were introduced in STK19: ΔRNAPII (D154A-T158A-V161A-N162A-V168A-W174A-W175A), ΔUVSSA (K203A-Y204A), and ΔCSA (R72A-T73A-D76A-D98A). Bars represent the means of 3 experiments with individual datapoints shown as colored circles. **i.** Endogenous RNAPII-S2 immunoprecipitation (IP) in RPE1 cells as in **b** to detect STK19-TY1 association with the TCR complex in parental and two *CSA-*KO clones.

### A cryo-EM structure of the RNAPII – TCR – ELOF1 – STK19 complex

To understand how STK19 integrates into the complex with arrested RNAPII, we bound purified STK19 to a reconstituted RNAPII-TCR complex comprised of the RNAPII elongation complex, CSB, CSA-DDB1, UVSSA, and ELOF1 (Kokic et al., 2021; Kokic et al., 2024). The resulting RNAPII-TCR-STK19 complex was isolated by size-exclusion chromatography and analyzed by cryogenic electron microscopy (Figure S5a-d). We obtained a structure of the complex at an overall resolution of 3.3 Å, which allowed for building a molecular model with good stereochemistry (Supplementary Table 7). We resolved all structured parts of STK19, comprised of three tightly packed winged-helix (WH) domains (Li et al., 2024), which we refer to as: the N-terminal domain 1 (residues 1-104), the middle domain 2 (residues 105-177) and the C-terminal domain 3 (residues 178-254) (Figure 4c,d).

STK19 directly interacts with RNAPII by docking domain 2 onto the RNAPII clamp head Zn-finger (Figure 4c,d,e). Domain 3 further extends towards and grips the downstream DNA by a high-density cluster of positively charged residues, thereby, in effect, extending the RNAPII DNA-binding clamp. Domain 3 further inserts a beta hairpin, with residues K203 and Y204 at the tip, into a pocket formed by the first three helices of UVSSA’s VHS domain. Interestingly, this interaction results in UVSSA movement toward the downstream DNA, which is the only major conformational change detected in the bound TCR factors as induced by STK19 binding (Figure 4c,d,f) (Kokic et al., 2024). Domain 1 of STK19 forms an extensive interface with blades 4, 5 and 6 of the CSA beta propeller, which likely further stabilizes the position of STK19 within the TCR complex (Figure 4c,d,g).

### The interplay of STK19 with other TCR factors regulates repair

To investigate whether the observed interactions of STK19 in the RNAPII-bound TCR complex have functional relevance, we generated a number of STK19 point-mutants predicted to perturb specific protein-protein interactions within the TCR complex (Figure 4e,f,g, Figure S6a). All mutant STK19 proteins were stably expressed and localized to the cell nucleus (Figure S6b,c). The point-mutants were designed to attenuate interactions with RNAPII (named ΔRNAPII), CSA (ΔCSA), and UVSSA (ΔUVSSA), respectively. Clonogenic Illudin S survival assays showed that the ΔRNAPII and ΔCSA mutants were as sensitive as *STK19*-deficient cells, whereas ΔUVSSA showed an intermediate phenotype compared to cells rescued with wild-type STK19 (Figure 4h). These point-mutants also failed to rescue TCR-dependent trabectedin incisions, while wild-type STK19 included in parallel did rescue (Figure S6d). Intriguingly, in spite of their phenotypic consequence, co-IP of endogenous RNAPII showed that each of the STK19 mutants could still be incorporated into the TCR complex after UV irradiation (Figure S6e) This is important as it suggests that multiple independent interactions contribute to STK19’s association with the TCR complex, with each individual interaction being necessary for proper positioning and function of the TCR factors within the complex, but not for the establishment or overall stability of the complex.

We note that knockout of *CSA*, which also impairs UVSSA recruitment (van der Weegen et al., 2020), abolished STK19 incorporation into the TCR complex (Figure 4i). In summary, STK19 wedges between elongating RNAPII, UVSSA, and CSA, opening the possibility that arrested RNAPII decorated with TCR factors, particularly CSA and UVSSA, constitutes the platform for STK19 recruitment, which is in turn crucial for the correct functioning of the complex. Moreover, STK19 positioning above the downstream DNA ideally places this TCR factor to mediate interactions with the incoming DNA repair machinery, with TFIIH likely accessing the DNA lesion from this direction (He et al., 2016; Kim et al., 2023; Kokic et al., 2019; Schilbach et al., 2021).

### STK19 positions and stabilizes TFIIH in the RNAPII-bound TCR complex

The location of STK19 in the TCR complex uniquely positions it to interact with TFIIH, which is located in front of RNAPII during transcription initiation (Aibara et al., 2021; He et al., 2016). To investigate this possibility, we employed AlphaFold (Jumper et al., 2021) to explore potential interactions between individual TFIIH subunits or the entire TFIIH complex and STK19. This analysis resulted in a high confidence score only for STK19 interacting with the repair helicase XPD (Figure 5a). The interaction surface on STK19 partially overlaps with the binding of UVSSA centered around Y204, with additional contacts that are unique for the STK19-XPD interaction (Figure 5a, b). Importantly, the obtained STK19-XPD model could subsequently be superimposed together with that of the TFIIH core complex (Kokic et al., 2019) onto the RNAPII-bound TCR complex without any steric clashes (Figure 5c,d). In this configuration, both XPB and XPD can engage the downstream DNA below STK19 (Figure 5c). These findings support a model in which STK19 positions and stabilizes TFIIH in the RNAPII-bound TCR complex in an optimal configuration for XPD to start scanning the transcribed strand for the DNA lesion (Kim et al., 2023). To analyze the potential functional consequences of STK19-TFIIH interaction, we monitored the 5′–3′ XPD helicase activity of the recombinant core TFIIH complex on a double-stranded DNA substrate (Kim et al., 2022; Kokic et al., 2019). TFIIH showed basal 5′–3′ helicase activity, which was weakly stimulated by recombinant STK19 protein (Figure 5e). These experiments suggest that STK19 may stabilize and potentially even stimulate TFIIH during transcription-coupled DNA repair. The overlap in binding sites suggests that UVSSA may be displaced once XPD is stabilized by docking onto RNAPII-bound STK19 (Figure 5a,b). In apparent agreement with this idea, immunoprecipitation of TCR complexes in a time-course revealed decreasing binding of UVSSA over time in WT, but persistent binding in *STK19*-KO cells, accompanying the less stable binding of TFIIH to RNAPII at all time-points after UV exposure (Figure 5f).

**Figure 5.**
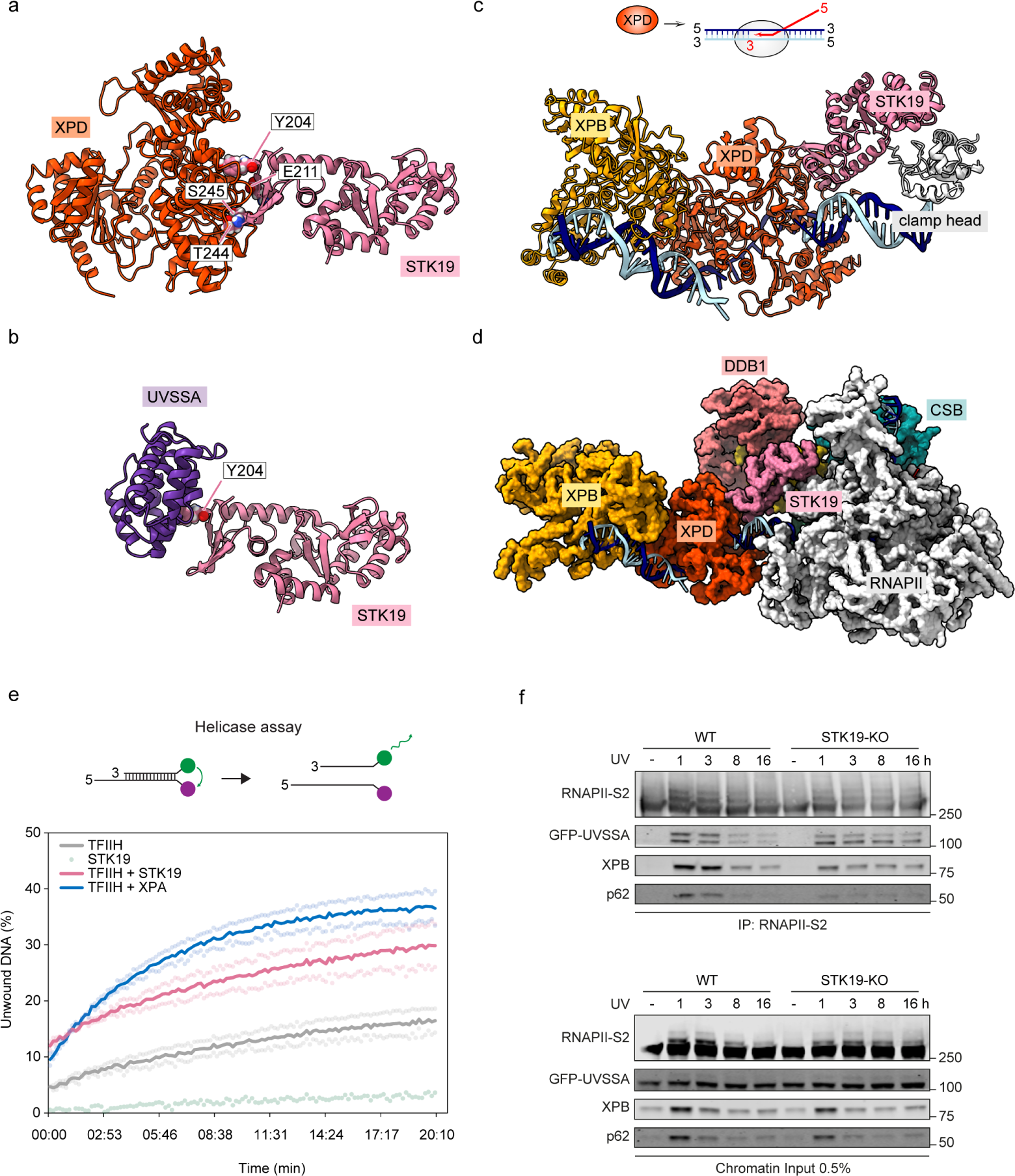
STK19 may position TFIIH in the RNAPII-TCR complex. **a.** AlphaFold model of an interaction between STK19 and XPD. **b.** Interaction interface between STK19 and UVSSA. **c.** The XPD-STK19 model with XPB-XPD bound to DNA from (Kokic et al., 2019) was superimposed on RNAPII-bound STK19. **d.** As in **c** now also showing the RNAPII-TCR and TFIIH core complexes. **e.** Schematic of the in vitro XPD helicase assay and real-time fluorescence measurement of the XPD 5′–3′ helicase activity using a fluorescence energy transfer-based assay when TFIIH (0.4 µM), STK19 (2.4 µM), or TFIIH (0.4 µM) + STK19 (2.4 µM), or TFIIH (0.4 µM) + XPA (1.2 µM) were added to the fluorescent substrate. **f.** Endogenous RNAPII-S2 immunoprecipitation (IP) in RPE1 cells with the indicated genotype that were untreated or at 0 h, 3 h, 8 h, or 16 h following UV-C irradiation (12 J/m^2^). TCR complex assembly was analyzed with the indicated antibodies. The data shown represent at least three independent experiments.

### STK19 suppresses the genome-wide accumulation of stalled RNAPII

The results above indicate that STK19 interacts with lesion-stalled RNAPII while having little or no impact on the assembly of the TCR complex itself. However, STK19 is crucial for the subsequent TCR process, including dual incision and the DNA synthesis necessary for gap filling. We reasoned that STK19 might affect RNAPII removal to allow the downstream repair reaction. If, for example, RNAPII were not removed efficiently from sites of DNA damage in the absence of STK19, this could interfere with the recruitment of downstream repair proteins to the DNA lesion, which would be shielded by stalled RNAPII (Nieto Moreno et al., 2023).

To detect persistent stalling of RNAPII in a genome-wide manner, we first employed strand-specific ChIP-seq using antibodies against elongating, Ser2-phosphorylated RNAPII, known as TCR-seq. The DNA fragments that co-purify with RNAPII from UV-irradiated cells are enriched for UV-induced photoproducts specifically in the transcribed stand (Nakazawa et al., 2020; van der Weegen et al., 2021). Strand-specific PCR amplification during library preparation therefore preferentially amplifies fragments from the non-damaged coding strand, resulting in a significant reduction in reads originating from the transcribed strand containing the DNA lesion (Figure 6a). The resulting strand bias after UV is therefore a direct consequence of RNAPII stalling at DNA lesions (Nakazawa et al., 2020).

**Figure 6.**
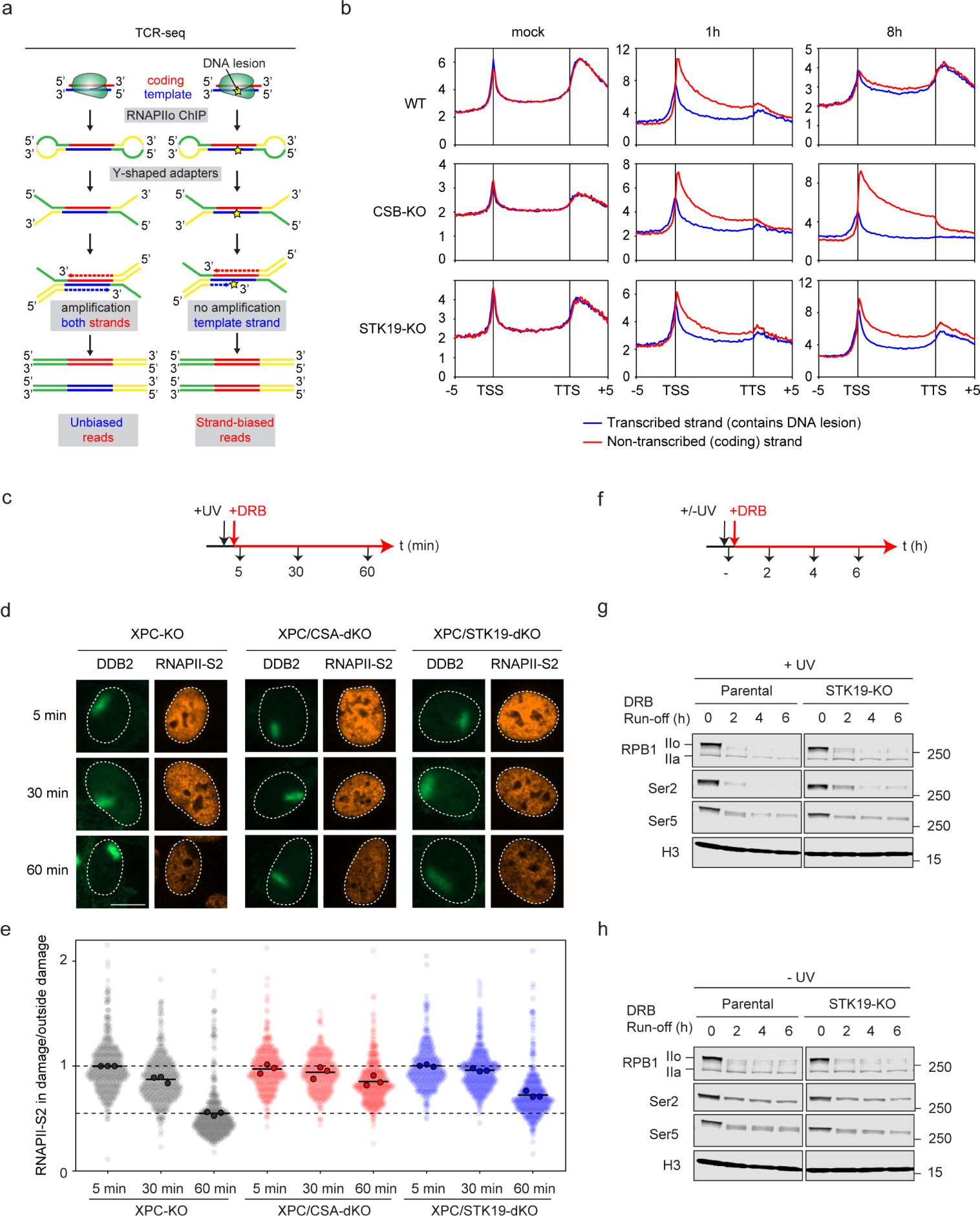
STK19 accelerates RNAPII clearance from sites of DNA damage. **a.** Schematic representation of TCR-seq. DNA that is co-purified with RNAPII after ChIP from UV-irradiated cells is highly enriched for DNA lesions in the transcribed strand, causing a strand-bias during library preparation (obtained from (van der Weegen et al., 2021)) **b.** Averaged metaplots of Ser2-RNAPII TCR-seq of 3,000 genes from the TSS until the TTS (-5 kb, +5 kb respectively) in RPE1 cells with the indicated genotype after mock-treatment or at 1 h, or 8 h after UV irradiation (12 J/m^2^). The coding (non-transcribed) strand is shown in red, while the template (transcribed) strand is shown in blue. **c.** Outline of a DRB run-off imaging assay. **d.** Representative images of RNAPII-S2 signal in RPE1 cells deficient in GGR (*XPC-*KO cells) with the indicated genotype at 5 min, 30 min, and 60 min after local UV damage (100 J/m^2^) marked by DDB2. Cells were treated with DRB immediately after UV irradiation to prevent new RNAPII molecules from moving into gene bodies. Scale bar, 10 μm. **e.** Quantification of **d**. The RNAPII-S2 mean pixel intensity at sites of local damage was divided by the mean pixel intensity outside local damage in the same cell nucleus to correct for a decrease of RNAPII-S2 signal due to a general RNAPII run-off caused by DRB. A ratio below 1 signifies that RNAPII-S2 loss in the local damage is faster than the general run-off outside the local damage. All cells are depicted as individual semi-transparent data points with the bar representing the mean of all data points. The individual means of three biological replicates are depicted as solid circles with black lines. **f.** Outline of a DRB-run off western blot assay. **g.** Western blot analysis on HEK293 cells using the experimental approach in **f**. Samples collected at the indicated time points after DRB addition immediately after UV irradiation. The abundance of total RPB1 (IIo – phosphorylated and IIa-non phosphorylated), S2-phosphorylated and S5-phosphorylated RPB1, as well as histone H3 (control) in chromatin is shown. **h.** Western blot analysis of experimental approach in **f**. DRB added without UV irradiation, and samples collected at the indicated time points. The abundance of total RPB1 (IIo – phosphorylated and IIa-non phosphorylated), S2-phosphorylated and S5-phosphorylated RPB1, as well as histone H3 (control) in chromatin is shown.

Capturing DNA fragments bound by RNAPII using this TCR-seq protocol showed no difference between WT, *STK19*-deficient, or *CSB*-deficient cells that had not been UV-irradiated (Figure 6a,b; mock). Metagene profiles of 3,000 genes revealed a distinct RNAPII distribution pattern, characterized by an enrichment of RNAPII at the transcription start site (TSS) and transcription termination site (TTS), respectively. Additionally, this analysis indicated that a similar number of sequencing reads originated from the transcribed and coding strands of the genes (Figure 6b). At 1 h after UV irradiation, however, we detected a strong strand bias in all cell-lines, which was completely resolved in WT cells within the initial 8 h following UV irradiation. By contrast, both *STK19*-deficient and *CSB*-deficient cells still showed a strand bias at 8 h after UV, indicating that stalled RNAPII complexes persist in the genomes of these cells.

### STK19 stimulates timely removal of DNA damage-stalled RNAPII

The persistence of damage-stalled RNAPII in *CSB*-KO cells above is due to the failure to assemble a TCR complex, but the result in *STK19*-KO cells suggests that although TCR complexes are formed in these cells, the obligatory removal of RNAPII does not take place. The detected RNAPII stalling in our TCR-seq experiments could reflect recurrent stalling of new polymerases at the same lesion, or persistent stalling of the same polymerase.

To investigate this further, we used two complementary methods in two different genetic backgrounds. First, we developed an imaging-based assay to investigate the fate of RNAPII at sites of DNA damage in RPE1 cells. To this end, we again used *XPC-*KO cells to exclude potential interference by GGR, and we locally irradiated cells with UV-C light through 5 µm pores (Moné et al., 2001). Immediately after UV irradiation, we treated cells with transcription elongation inhibitor 5,6-dichloro-1-β-D-ribofuranosylbenzimidazole (DRB) to prevent new RNAPII complexes from transitioning from initiation to elongation, enabling us to specifically monitor the fate of already elongating RNAPII complexes (Figure 6c). In the absence of DNA damage, immunofluorescence staining of elongating RNAPII after DRB treatment to inhibit new transcription showed a gradual reduction in the general nuclear signal for Ser2-RNAPII within a time-course of 60 min, consistent with RNAPII elongating during that time and eventually running off even long genes and becoming de-phosphorylated (Figure 6d, *XPC*-KO panels). However, at sites of local UV damage, marked by staining for DDB2, we measured an accelerated loss (∼2-fold) of RNAPII signal compared to elsewhere in the nucleus (Figure 6e, *XPC-*KO). This loss reflects the active removal of DNA damage-stalled RNAPII from chromatin. Indeed, knockout of *CSA* strongly decreased the accelerated loss of RNAPII from sites of DNA damage, while *STK19*-deficient cells showed slower and attenuated RNAPII removal compared to WT cells (Figure 6d,e). A similar attenuated clearance of RNAPII from sites of DNA damage was observed in single *STK19*-KO cells in a GGR-proficient genetic background (Figure S6f,g).

As a complementary approach, we used western blot analysis to monitor the different RNAPII forms in chromatin after DRB treatment in HEK293 cells (Figure 6f). Following UV irradiation, a marked delay in Ser2-RNAPII (marking the elongating form) run-off was detected in *STK19*-deficient cells compared to WT cells. We note that the Ser2-phosphorylated, elongating form of RNAPII was the main form that persisted in *STK19*-deficient cells, while no clear differences were observed for the Ser5-phosphorylated, promoter-proximal RNAPII (Figure 6g). As a control, such differences between WT or *STK19*-deficient cells were not observed in the absence of DNA damage (Figure 6h). Together, these data indicate that STK19 promotes the timely removal of DNA damage-stalled RNAPII from TCR complexes in chromatin, required to enable the downstream DNA repair steps.

### STK19 is required for the timely clearance of ubiquitylated RNAPII

The results on RPB1 turnover after DNA damage shown above are important as both CSB and CSA are recruited normally in *STK19-*KO cells, so that ubiquitylation of damage-stalled RPB1 would be expected to occur normally. To investigate the impact of STK19 on RNAPII ubiquitylation in more detail, we performed pull-down experiments using GST-DSK2, enabling the enrichment of native, poly-ubiquitylated proteins (Anindya et al., 2007). As expected, DSK2 pull-down from HEK293 cell extracts followed by detection of total RNAPII or elongating Ser2-RNAPII revealed a dramatic increase in UV-induced ubiquitylation 45 min after UV irradiation, which decreased at 3 h and completely reverted to background levels within 24 h after UV irradiation in WT cells (Figure 7a). By contrast, persistent ubiquitylated RNAPII on chromatin was detected in *STK19*-deficient cells at both 3 h and 24 h after UV irradiation (Figure 7a). To further investigate the persistence of ubiquitylated RNAPII, we performed immunoprecipitation of TCR complexes in a time-course after UV irradiation in RPE cells. In the WT, RNAPII ubiquitylation and binding of TCR proteins to RNAPII was prominently observed at early time-points (3 h), which gradually decreased up to 16 h after UV (Figure 7b). In contrast, while *STK19*-KO cells initially displayed normal RNAPII ubiquitylation and association of TCR proteins, this was more persistent 8 h after UV (Figure 7b). This suggests that while TCR complex assembly is normal even in the absence of STK19, the RNAPII-bound TCR complexes are not disassembled normally without it.

**Figure 7.**
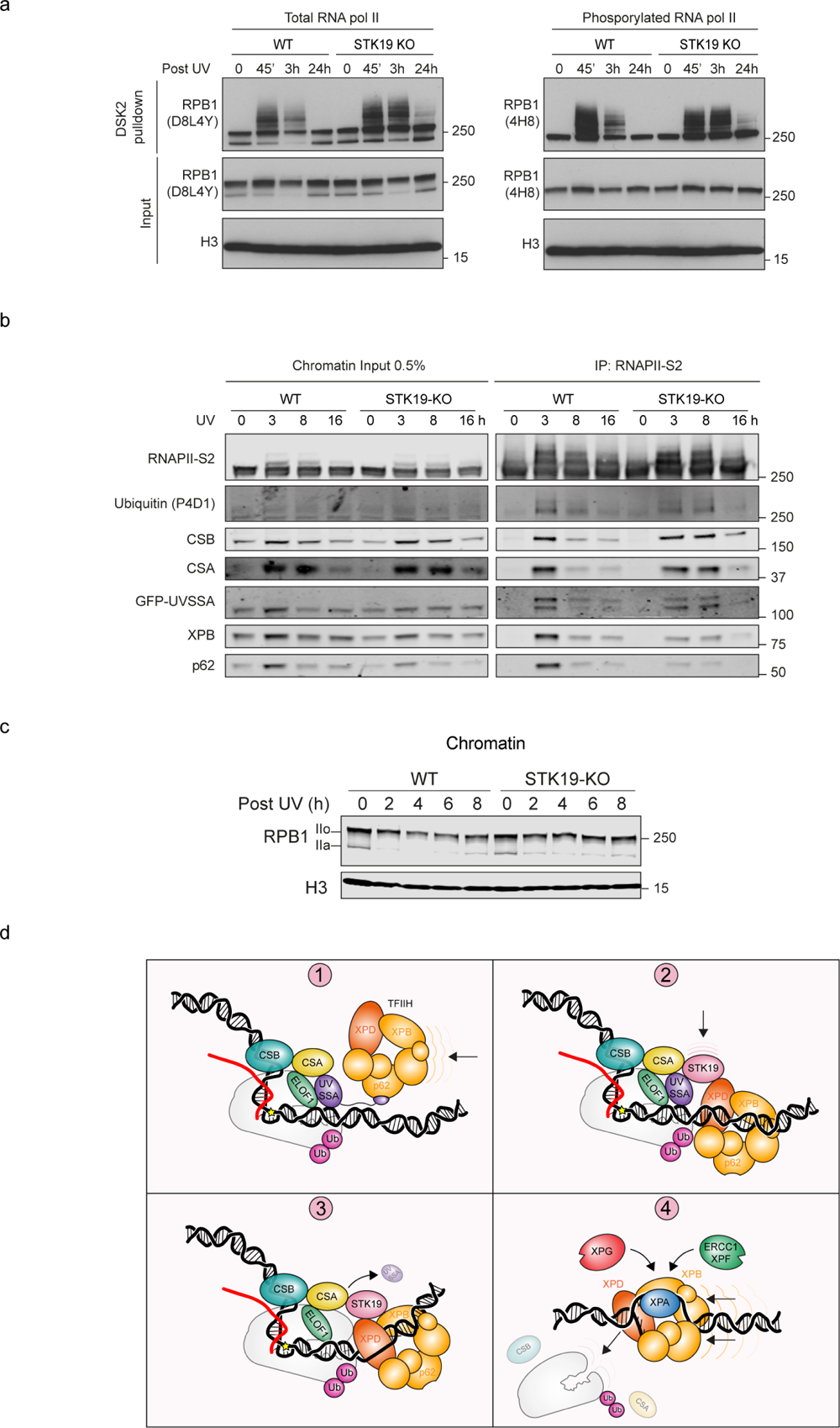
Ubiquitylated RNAPII persists on chromatin when STK19 is absent. **a.** Dsk2 pulldown-western blot analysis of HEK293 cells with the indicated genotype that were untreated or at 45 min, 3 h, or 24 h following UV-C irradiation (15 J/m^2^). RNAPII was detected with the indicated antibodies. **b.** Endogenous RNAPII-S2 immunoprecipitation (IP) in RPE1 cells with the indicated genotype that were untreated or at 0 h, 3 h, 8 h, or 16 h following UV-C irradiation (12 J/m^2^). TCR complex assembly and RNAPII ubiquitylation were analyzed with the indicated antibodies. The data shown represent at least three independent experiments. **c.** Western blot of a time course after UV irradiation showing the amount of unphosphorylated (IIa) and phosphorylated (IIo) RPB1 on chromatin in WT and STK19-KO Flp-In T-REx HEK293. H3 is used as loading control. **d.** Molecular model for how STK19 accelerates effective clearance of arrested RNAPII to facilitate transcription-coupled DNA repair. (1) TCR complex assembly with TFIIH recruitment by UVSSA through an interaction with p62. (2) STK19 wedges between CSA, UVSSA, and RNAPII, and positions TFIIH through an interaction with XPD. (3) UVSSA is displaced and XPD unwinds towards RNAPII. (4) TFIIH displaces RNAPII, providing downstream NER factors access to the lesion.

The finding that RNAPII is cleared more slowly from chromatin after DNA damage, yet seems to be more ubiquitylated, would be consistent with a model in which proteolysis of ubiquitylated RNAPII occurs more slowly. Indeed, degradation of RPB1 was somewhat delayed in *STK19*-KO cells, both in HEK293 (Figure 7c) and RPE1 cells (Figure S7a,b), while over-expression of STK19 appeared to stimulate UV-induced RPB1 degradation (Figure S7c). Together, these findings are consistent with a model in which, in the absence of STK19, the early TCR factors remain associated with RNAPII, which is ubiquitylated normally but not displaced in a timely manner from chromatin, thereby interfering with the downstream repair steps (Figure 7d).

## Discussion

While the basic mechanisms of most DNA repair processes have now been uncovered, much less is understood about the factors and molecular events governing TCR. For example, although the initial recruitment of TCR proteins to damage-stalled RNAPII has been described, it is still unclear how TCR transitions to downstream repair steps, allowing removal of RNAPII and access to the DNA lesion. The work described here identifies STK19 as a new TCR factor that promotes this transition. Indeed, while initial TCR complex assembly and RNAPII ubiquitylation are not affected or decreased by the loss of STK19, timely clearance of lesion-stalled RNAPII is impaired, ultimately interfering with downstream repair steps. Cryo-EM analysis revealed that STK19 binds to the RNAPII-bound TCR complex, locating it between RNAPII, UVSSA, CSA, and above the DNA in front of lesion-arrested RNAPII. This ideally positions STK19 to mediate interactions with incoming DNA repair proteins. Indeed, integrating our cryo-EM structure with a previous TFIIH structure and AlphaFold-based molecular modeling supports a model in which STK19 plays a role in positioning the ATPase subunits of TFIIH onto the downstream DNA.

### STK19 is not a kinase and functions specifically in TCR

We previously reported that the start site of the *STK19* coding region is incorrectly annotated in the human genome (Rodriguez-Martinez et al., 2020). Despite its name suggesting a kinase of 364 amino acids, the *STK19* gene actually encodes a 254-amino acid protein, the function of which was previously unknown except for a poorly defined role in the DNA damage response (Boeing et al., 2016; Olivieri et al., 2020). Here, we demonstrate that an sgRNA targeting the mis-annotated exon immediately upstream of the true start codon does not affect STK19 function and thus does not induce Illudin S sensitivity. Conversely, an sgRNA targeting the subsequent exon, which is the first according to the revised annotation, triggers all the TCR phenotypes observed here. Theoretically, both the long and a short isoform of STK19 would be knocked out by this approach. However, all the phenotypes were rescued by re-expressing the 254-amino acid cDNA of STK19, demonstrating that this isoform of STK19 contains all functions required for TCR. The detailed analysis of STK19 function provided here reveals that STK19 is crucial for TCR, without affecting global genome NER, or the response to replication stress and DNA-double strand breaks.

That STK19 is not a kinase also became abundantly clear through the recently published crystal structure of STK19 which does not show evidence of a kinase domain (Li et al., 2024). The crystal structure of STK19 alone matches that observed in our cryo-EM structure of the entire TCR complex (Figure S7d). Notably, (Li et al., 2024) observe that STK19 can bind DNA via a cluster of positively charged residues (K186, K190, K201). This aligns with our cryo-EM structure, which shows that this STK19 cluster interacts with the DNA downstream of RNAPII in the TCR complex. Intriguingly, Li et al. suggested that STK19 binds DNA as a homodimer (Li et al., 2024), while our structure reveals that STK19 is incorporated into the TCR complex as a monomer. Indeed, a dimeric STK19 would be incompatible with our structure, as we observe extensive interactions with UVSSA (and potentially XPD) at the dimerization interface mapped by (Li et al., 2024). Nonetheless, functions of dimeric STK19 outside TCR cannot be excluded.

### STK19 regulates the positioning of TFIIH during TCR

Cells that are defective in CSB, CSA, UVSSA, ELOF1, or RNAPII ubiquitylation show strongly impaired recruitment of TFIIH to DNA damage-stalled RNAPII (Kokic et al., 2024; Nakazawa et al., 2020; van der Weegen et al., 2021; van der Weegen et al., 2020), which helps explain the repair defect displayed by these cells. In contrast, despite TCR being largely inactivated in cells lacking STK19, its absence only marginally affects TFIIH recruitment to stalled RNAPII. Previous work established that TFIIH is initially recruited to the TCR complex by UVSSA (Fei and Chen, 2012; van der Weegen et al., 2020) through the p62 subunit of TFIIH (Okuda et al., 2017). We suggest that STK19 correctly positions the TFIIH ATPase subunits XPB and XPD onto the DNA in front of the lesion-stalled RNAPII (Figure 7d). This is conceptually reminiscent of the role proposed for ELOF1 in TCR complex assembly, which is to ensure the correct positioning of CSA after its recruitment by CSB (Kokic et al., 2024). Due to its numerous interactions, STK19 could therefore be considered a ’molecular glue’ that stabilizes and correctly positions various proteins within the TCR complex. We therefore interpret the attenuated binding of TFIIH in the absence of STK19 as incorrectly positioned and therefore less stably bound TFIIH, which is more easily lost in our IP procedure. It is worth noting that although *STK19* KO clearly dramatically affects TCR and cellular sensitivity to different DNA damage-inducing treatments, the effects are not as dramatic as those observed in *CSB*-KO cells. Whether STK19 is more important for certain types of transcription-blocking lesion than for others, or if CSB mutation also affects the response to DNA-damaging treatments via other repair pathways than the classical TCR pathway, remains to be established.

In addition to the interactions observed in our cryo-EM structure, which revealed that STK19 wedges between RNAPII, CSA, and UVSSA, we utilized AlphaFold to explore interactions with downstream repair proteins that were missing from our structure (Jumper et al., 2021). This approach yielded a high-confidence interaction between STK19 and the XPD repair helicase subunit of TFIIH. This model enabled us to superimpose the DNA-bound TFIIH core complex (Kokic et al., 2019) onto the cryo-EM structure of the RNAPII-bound TCR complex including STK19, providing a compelling glimpse of what this complex might look like. Remarkably, the trajectory of the DNA engaged by XPB and XPD from the previous structure aligns with the DNA in front of RNAPII bound by STK19, suggesting that STK19 is ideally positioned to guide both TFIIH ATPases onto the DNA in front of lesion-arrested RNAPII. Obtaining a cryo-EM structure of a lesion-arrested RNAPII complex bound by TCR factors, including TFIIH, is an important future goal. However, this will require in vitro reconstitution of TCR, which has not so far been possible.

### The fate of DNA damage-stalled RNAPII

Different models have been proposed to explain what happens to RNAPII after it stalls on a lesion in the DNA. For example, it was suggested that RNAPII backtracks to allow NER proteins access to the DNA lesion, enabling the polymerase to continue after DNA repair (Tornaletti et al., 1999). Here, with the use of a new assay, we present evidence indicating that Ser2-phosphorylated, elongating RNAPII disappears from damage sites following local UV treatment. This supports the notion that although RNAPII backtracking may occur, it is likely succeeded by RNAPII removal, and subsequently, the polymerase does not continue transcription at the site of DNA repair. In support of this notion, previous measurements of genome-wide nascent transcription and RNAPII occupancy also suggest that transcription primarily restarts from gene promoters following UV-induced transcriptional arrest (Andrade-Lima et al., 2015; Chiou et al., 2018; Tufegdzic Vidakovic et al., 2020; van den Heuvel et al., 2021a).

Another model proposes that lesion-stalled RNAPII undergoes ubiquitylation, is extracted from chromatin, and is degraded by the proteasome as part of the ’last resort’ pathway (Nieto Moreno et al., 2023; Wilson et al., 2013). Indeed, *CSB*-KO and *CSA*-KO cell lines fail to efficiently ubiquitylate and degrade RNAPII in response to UV damage (Anindya et al., 2007; Bregman et al., 1996; Kokic et al., 2024; Nakazawa et al., 2020; Nakazawa et al., 2012). Notably, UVSSA-deficient cells also exhibit reduced RNAPII ubiquitylation, yet they degrade RNAPII normally in response to UV damage (Kokic et al., 2024; Nakazawa et al., 2020; Nakazawa et al., 2012). Conversely, *STK19*-KO cells, in which TCR-mediated ubiquitylation of RNAPII is unaffected and even prolonged, experience delayed RNAPII degradation. This leads us to hypothesize that ubiquitylation of RNAPII alone may not suffice for its properly regulated removal and degradation.

Our cryo-EM structure and AlphaFold modeling suggest that STK19 positions the TFIIH complex onto the DNA in front of lesion-arrested RNA(Laine and Egly, 2006)PII, transforming TFIIH from a transcription-regulatory factor into a DNA repair factor. Recent research has shown that XPD scans in the 5’ to 3’ direction along the lesion-containing strand (Li et al., 2015), which corresponds to the transcribed strand during TCR. Once it encounters the lesion and is inhibited by it, XPB pulls the DNA in the opposite direction, enlarging the DNA bubble for dual incision (Kim et al., 2023). We speculate that the coordinated actions of the TFIIH motors may destabilize and potentially even trigger the dissociation of RNAPII, which would be consistent with results obtained in a minimal RNAPII displacement system in vitro (Laine and Egly, 2006; Sarker et al., 2005). It is an intriguing possibility that STK19 serves as the bridge between initial TCR complex formation and downstream repair by enabling efficient TFIIH-mediated remodeling and ultimately displacing arrested RNAPII(Sarker et al., 2005).

Altogether, our findings collectively unveil the molecular mechanism through which STK19 directs TCR from transcription to DNA repair. It achieves this by reorganizing the TCR complex to facilitate TFIIH positioning onto the downstream DNA, culminating in RNAPII clearance, thereby providing downstream NER proteins access to the DNA lesion.

## Acknowledgments

MSL laboratory was supported by the Netherlands Scientific Organization (ENW grant OCENW.M20.056, and VIDI grant ALW.016.161.320), and an European Research Council (ERC) Consolidator Grant STOP-FIX-GO (grant agreement 101043815). JS laboratory was supported by the Novo Nordisk Foundation (Laureate grant NNF19OC0055875), the Danish National Research Foundation (Chair grant DNRF153, and Center of Excellence grant DNRF166), and by an ERC Advanced Grant TRANSDAM (Grant agreement 693327), and by an EMBO Fellowship (ALTF 911-2020) to NNM. JS was also funded by the Francis Crick Institute, which receives funding from Cancer Research UK [FC001166], the UK Medical Research Council [FC001166], and the Wellcome Trust [FC001166]. PC laboratory was supported by the Deutsche Forschungsgemeinschaft (SFB860, SPP1935), an ERC Advanced Grant TRANSREGULON (grant agreement No 693023), and the Volkswagen Foundation. ML laboratory was supported by NHGRI (5UM1HG009382) and NCI (5R01CA213214). ODS laboratory was supported by Korean Institute for Basic Science (IBS-R022-A1) and the NCI (US) (P01CA092584). The funders had no role in study design, data collection and analysis, decision to publish or preparation of the manuscript. We would also like thank Chi-Lin Tsai and John A. Tainer (MD Anderson, United States) for providing purified TFIIH protein,

## Data Availability

The electron density reconstructions and structure coordinates were deposited to the Electron Microscopy Database (EMDB) and to the PDB under the following accession codes: EMD-19909 and PDB 9ER2 for the RNAPII-TCR-ELOF1-STK19 structure.

## Author contribution

DvdH generated *STK19-*KO and STK19/*CSA-*dKO (RPE1), STK19 mutants and rescue lines, illudin S survival, trabectedin survival, RRS, TCR-seq & analysis, BrU-seq & analysis, RNAPII-IPs, ATF3 western blot analysis, wrote the paper. MR-M performed *STK19-*KO validations in Flp-In T-REx HEK293, Flp-In T-REx HeLa and MRC5VA. Survival and growth assays after UV, CPT, HU, NCS. Performed TT_chem_-seq. Generation of STK19 overexpression cell lines. DRB run-off experiments and RPB1 stability experiments. RPB1 immunoprecipitation. RNA extraction and qPCRs. Assisted with writing the manuscript. PJvdM generated rescue line in XPC/*STK19-*dKO cells, TCR-UDS, GGR-UDS, γH2AX trabectedin assay, BrU-seq, DRB run-off RNAPII imaging, RNAPII time-course IPs, wrote the paper. NNM performed DSK2 assay for detection of ubiquitylated RPB1, DRB-run off experiments and RPB1 stability experiments in HEK293 cells, RPB1 immunoprecipitation. JP performed comet assay after trabectedin. H-SK performed in vitro XPD helicase assays. JJMvS generated HMCES-KO and XRCC1-KO single and CSB-dKO cells, performed survivals, RRS assay and western blot analysis. APW generated XPC/*STK19-*dKO cells, performed RRS. AD’S performed modeling of the XPD-STK19 interaction. GY performed RPB1 stability experiments in RPE1 cells. AH performed DSK2 assay for detection of ubiquitylated RPB1. KK performed DSK2 assay for detection of ubiquitylated RPB1, RPB1 stability experiments, RPB1 immunoprecipitation. JW generated *STK19-*KO Flp-In T-REx HEK293, Flp-In T-REx HeLa and MRC5VA. TB performed survival and growth assays after DNA damage, asssistance with TCR-seq assays. RM performed TT_chem_-seq analysis. KA performed CRISPR screen on 4-NQO. KdL performed and analysed CRISPR screens on 4-NQO. IK performed network analysis of CRISPR screens. ML supervised RNA isolation and sequencing for BrU-seq. RMFW supervised CRISPR screen on 4-NQO. PC supervised the RNAPII-TCR-STK19 structure. ODS supervised comet assay after trabectedin and in vitro XPD helicase assay. GK purified TCR proteins, generated the RNAPII-TCR-STK19 cryo-EM structure, wrote the paper. JQS supervised the project, wrote the paper. MSL supervised the project, designed STK19 point-mutants, generated figures, wrote the paper. All authors commented on and approved the manuscript.

## Methods

### Cell lines

All cell lines are listed in Supplementary Table 1. Cell lines were cultured at 37°C in an atmosphere of 5% CO_2_ in DMEM GlutaMAX (Thermo Fisher Scientific) supplemented with penicillin/streptomycin (Sigma), and 8-10% fetal bovine serum (FBS; Bodinco BV or Thermo Fischer Scientific (Gibco)).

### Generation of knockout cells

Parental RPE1-hTERT cells stably expressing inducible Cas9 (iCas9) that are also knockout for *TP53* and the puromycin-N-acetyltransferase *PAC1* gene were described previously (referred to as RPE1-iCas9) (van der Weegen et al., 2021). RPE1-iCas9 cells were transfected with pU6-gRNA:PGK-puro-2A-tagBFP (Sigma–Aldrich library from the LUMC) containing 2 gRNAs against XPC, or with Cas9-2A-EGFP (pX458; Addgene #48138) containing a gRNA against CSA, ELOF1, STK19, HMCES or XRCC1 from the TKOv3 library using lipofectamine 2000 (Invitrogen). The sgRNAs are listed in Supplementary Table 2, and plasmids in Supplementary Table 3. Cells were FACS sorted on EGFP and plated at low density, after which individual clones were isolated, expanded, and verified by western blot analysis and/or Sanger sequencing using the primers listed in Supplementary Table 4. Indicated double knockout cell lines were subsequently generated in clonally selected single knockout cell lines.

CRISPR-Cas9-nuclease-mediated genome editing was performed in Flp-In T-REx HEK293, MRC5VA and Flp-In T-Rex HeLa cell lines. The oligonucleotide encoding the gRNA for targeting the coding region of *STK19* is described in Supplementary Table 2. The gRNA was annealed and ligated into pSpCas9(BB)-2A-GFP (Ran et al., 2013) (Addgene, PX458), and plasmids were sequenced after cloning and transformation. To generate knockouts, cells were transfected with pSpCas9(BB)-2A-GFP plasmids expressing the gRNA, EGFP and Cas9 using Lipofectamine 2000 (Thermo Fisher Scientific) according to the manufacturer’s instructions. 48 h after transfection, high GFP-positive cells were sorted clonally by fluorescence activated cell sorting (FACS) into 96-well plates and cultivated until colonies were obtained. For *STK19-*KOs, genomic PCRs around the edited site were sequenced and analyzed using the Web tool “TIDE” (https://tide.deskgen.com). Cells containing Indels were expanded from the master plate for further analysis by western blot. The expression of the upstream gene DXO was analyzed by RT qPCR to confirm that the mutation in *STK19* is not affecting the expression of the upstream gene. CSB-KOs were generated as described previously (Tufegdzic Vidakovic et al., 2020).

### Plasmids

All plasmids are listed in Supplementary Table 3. PGK-EGFP-C1-IRES-puro and PGK-EGFP-N1-IRES-puro plasmids were generated as described before (Kokic et al., 2024). EGFP in PGK-EGFP-N1-IRES-puro was replaced by a triple TY-tag to generate PKG-TY-IRES-puro or by 3xFlag-Bio-DDB2 to generate PGK-3xFlag-Bio-DDB2-IRES-Puro. The STK19 cDNA was amplified by PCR (Primers for cloning are listed in Supplementary Table 5) and inserted into pcDNA5 STK19-Flag-GFP, followed by subcloning into PKG-TY-IRES-puro. Three STK19 mutant cDNA constructs were generated by overlap PCR using primers described in Supplementary Table 5; PGK-STK19(154A-158A-161A-162A-168A-174A-175A)-TY-IRES-Puro (ΔRNAPII), PGK-STK19(72A-73A-76A-98A)-TY-IRES-puro (ΔCSA), and PGK-STK19(203A-204A)-TY-IRES-Puro (ΔUVSSA). The flag-tagged STK19 overexpression plasmid was constructed as previously described (Rodriguez-Martinez et al., 2020). Briefly, C-terminal 3xflag-tagged misannotated STK19 was generated by Genscript, and cloned into pFRT/TO (using EcoRV and XhoI sites). STK19 was generated by deleting the first 110 amino acids of the misannotated STK19 using Q5 site directed mutagenesis (NEB, E0554S) and addition of GFP was done using In-Fusion system (Takara 639649).

### Generation of stable cell lines

To visualize damaged DNA in *XPC-*KO RPE1-iCas9 cells, we transfected cells with PGK-3xFlag-Bio-DDB2-IRES-Puro using lipofectamine 2000 (Invitrogen), and selected on 1 µg/mL puromycin for at least 2 weeks to isolate stable polyclonal cell lines expressing flag-biotin-tagged DDB2. To stably express TY-tagged STK19 constructs (WT or mutant constructs), the indicated cell lines were transfected with PGK-STK19-TY-IRES-puro (WT, 154A-158A-161A-162A-168A-174A-175A (ΔRNAPII), 72A-73A-76A-98A (ΔCSA), 203A-204A (ΔUVSSA)) using lipofectamine 2000 (Invitrogen). Cells were subsequently selected on 1 µg/mL puromycin for at least 2 weeks to isolate polyclonal cell lines stably expressing TY-tagged STK19 protein. To enable better detection of UVSSA, RPE1-iCas9 *STK19-*KO (7) + STK19-TY(WT) cells were transfected with PGK-GFP-UVSSA-IRES-puro using lipofectamine 2000 (Invitrogen). Cells were selected on 1 µg/mL puromycin for at least 2 weeks and subsequently sorted on GFP to ensure homogenous expression. Homogeneous expression of all fusion proteins was confirmed by fluorescence microscopy and/or western blotting. Flp-In T-REx HEK293 cell lines expressing doxycycline inducible STK19-flag-GFP or STK19-3xflag were generated as described previously (Gregersen et al., 2014). Briefly, Flp-In T-REx HEK293 cell lines maintained in 100 μg/mL zeocin and 15 μg/mL blasticidin prior to transfection, were co-transfected with a 9:1 ratio of pOG44 Flp-recombinase expression vector (Thermo Fisher Scientific, V600520) and pFRT/TO/STK19-flag hygromycin resistant constructs using Lipofectamine 2000 (Thermo Fisher Scientific, 11668019) according to the manufacturer’s instructions. 24 h after transfection, cells were seeded as single cells and after another 24 h the cell culture media was supplemented with 100 μg/mL hygromycin (H011, TOKU-E) and 15 μg/mL blasticidin (B007, TOKU-E). Expression of tagged proteins was induced overnight by the addition of doxycycline (Clontech, 8634-1, 10 ng/mL final concentration) and all clones were verified by western blotting using antibodies against flag.

### Knockout validation by Sanger sequencing of genomic DNA

Genomic DNA was isolated by resuspending cell pellets in whole cell lysate (WCE) buffer (50 mM KCL, 10 mM Tris pH 8.0, 25 mM MgCl_2_, 0.1 mg/mL gelatin, 0.45 % Tween-20, 0.45 % NP-40) containing 0.1 mg/mL Proteinase K (EO0491;Thermo Fisher Scientific) and incubating for 2 h at 56°C followed by a 10 min heat inactivation of Proteinase K by 96°C. Samples were further purified by incubation with 30% v/v of 5 M NaCl for 5 min, followed by centrifugation at 20.000 g for 10 min and removal of protein-containing pellet. Purified gDNA was precipitated by addition of equal volumes of isopropanol, and DNA pellets were washed with 70% ethanol. Fragments of approximately 1-2 kb spanning the guide RNA target sites or the genomically integrated cDNA were PCR amplified using a Taq polymerase mastermix (Qiagen), followed by Sanger sequencing using the primers listed in Supplementary Table 4.

### Western blotting

Total cell lysates in RPE1-iCas9 cells were harvested by scraping cells in Laemmli-SDS sample buffer. Cell lysates or immunoprecipitation samples were boiled for 10 min at 95°C. Proteins were separated on Criterion™ XT Bis-Tris 4–12% Protein Gels (Biorad; 345-0124 or 345-0125) in MOPS Running Buffer (Thermo Fisher, NP0001-02). Then, blotted onto PVDF membranes (IPFL00010, EMD Millipore) in Tris/glycine blotting buffer (0.025 M Tris, 0.192 M glycine) with 20% methanol. Membranes were blocked with commercial blocking buffer (Rockland, MB-070-003) or with 5% fat-free milk in PBS with 0.1 % Tween-20 for 1 h at room temperature. Membranes were then probed with indicated antibodies in commercial blocking buffer or 5% fat-free milk in PBS with 0.1 % Tween-20 (Antibodies are listed in Supplementary Table 6). Proteins were stained with fluorochrome-conjugated secondary antibodies and were detected on an Odyssey CLx system and Image Studio software (Li-Cor). For whole cell extracts in FlpIn TRex HEK293, FlpIn TRex HeLa or MRC5VA, cell pellets were lysed in NP-40 lysis buffer (50 mM Tris-HCl pH 7.5, 500 mM NaCl, 2 mM EDTA, 0.5% (v/v) NP-40, 0.5 mM DTT, PhosSTOP (Sigma-Aldrich, 04906837001) and Protease Inhibitor Cocktail (Sigma-Aldrich, 05056489001). Chromatin fractionation was carried out as described previously (Gregersen et al., 2019). 30–100 μg protein/lane was separated on 3%–8% Tris-Acetate gels (BioRad, 3450130) and transferred to nitrocellulose membranes (GE Healthcare Life Sciences, 10600002). Membranes were blocked in 5% (w/v) skimmed milk in PBS-T (PBS, 0.1% (v/v) Tween20) for 1 h at room temperature and incubated with primary antibody (in 5% (w/v) skimmed milk in PBS-T) overnight at 4°C. Primary antibodies are listed in Supplementary Table 6. Antibody against vinculin or histone H3 were used to control loading. Membranes were washed several times in PBS-T, incubated with HRP-conjugated secondary antibody in 5% (w/v) skimmed milk in PBS-T and visualized using SuperSignal West Pico PLUS (for Vinculin and H3), Dura (for flag and RPB1) Chemiluminescent Substrate ECL reagent (Thermo Fisher Scientific, 34577 or 34075) or Radiance Plus Femtogram HRP substrate (for endogenous STK19) (Azure Biosystems, AC2103).

### Clonogenic survival assays

RPE1-iCas9 cells were plated in low density in medium containing indicated concentrations of Illudin S (Santa Cruz, sc-391575) or Trabectedin (Medchem express, HY-50936). Cells were allowed to form clones for 9–10 days in the presence of Illudin S or Trabectedin. To visualize clones, cells were subjected to NaCl fixation and methylene blue staining. Flp-In T-REx HEK293 cells were seeded into 6-well plates and treated with the indicated doses of the specified drugs or UV-C. Colonies were fixed by 4% (v/v) formaldehyde 11 days after seeding and stained with a 0.1% (w/v) crystal violet solution. Colonies from 3 biological replicates were counted. Cell survival after treatment was defined as the percentage of cells able to form clones, relative to the untreated condition.

### Sensitivity assays

For growth analysis using Incucyte (Sartorius) cells where treated with the indicated drug concentration for the indicated time or with UV-C using a custom built conveyor-belt and monitoring the exact dose using a UV-meter (Progen Scientific). Analysis were performed in 96-well plates with starting seeding densities of 10-20%. Luminescence growth assays (camptothecin (CPT), Hydroxyurea (HU) and neocarzinostatin (NCS)), were performed using CellTiter-Glo Luminescent Cell Viability Assay (Promega G7570) following manufacturer instructions.

### Immunoprecipitation of endogenous RNAPII

Cells were mock treated or irradiated with UV-C light (12 J/m^2^) and harvested at indicated timepoints after UV. For endogenous RNAPII-S2 IPs, chromatin-enriched fractions were prepared by lysing the cells for 20 min on a rotating wheel at 4° C in 1 mL EBC-1 buffer (50 mM Tris [pH 7.5], 150 mM NaCl, 2 mM MgCl_2_, 0.5% NP-40, and protease inhibitor cocktail (Roche)), followed by centrifugation, and removal of the supernatant. The chromatin-enriched cell pellets were resuspended in 1 mL ECB-1 buffer supplemented with 500 U/mL Benzonase® Nuclease (Novagen) and 2 µg RNAPII-S2 (ab5095, Abcam) for 1 h at 4 °C. The salt concentration was subsequently increased to 300 mM NaCl by addition of 5 M NaCl, and the samples were incubated for another 30 min on a rotating wheel at 4 °C. Samples were then centrifuged for 10 min at 14.000 rpm at 4 °C. 50 µL of the supernatants were saved as input fraction and lysed in 50 µl Laemmli-SDS sample buffer. The rest of the sample was transferred to fresh tubes. The protein complexes were immunoprecipitated by incubation with 20 µL prewashed Protein A agarose beads (Millipore) for 90 min at 4°C. After incubation, the beads were washed 6 times with ECB-2(300) buffer (50 mM Tris [pH 7.5], 300 mM NaCl, 1 mM EDTA, 0.5% NP-40, and protease inhibitor cocktail (Roche)). After final washes, beads were resuspended in 20 µl Laemmli-SDS sample buffer. For subsequent analysis by western blotting, the samples were boiled for 10 min at 95°C.

### Immunoprecipitations of STK19

Flp-In T-REx HEK293 cells stably expressing doxycycline (Dox)-inducible flag-STK19 were induced overnight by the addition of Dox (10 ng/mL final concentration). Cells were harvested by scraping in ice-cold PBS, washed once in cold PBS and pelleted by centrifugation at 1,500 rpm for 5 min at 4°C. Cells were then fractionated as previously described and all chromatin fractions were pooled to enrich in STK19 and RPB1. Phosphatase inhibitors (PhosSTOP, Sigma-Aldrich, 04906837001) and Protease Inhibitor Cocktail (Sigma-Aldrich, 05056489001) were added fresh to all buffers. Flag immunoprecipitation was done by incubating chromatin fractions with ANTI-FLAG M2 Affinity Gel (Sigma-Aldrich, A2220) at 4°C for 3 h. Beads were washed 5 times in IP wash buffer (150 mM NaCl, 20 mM Tris-HCl pH 7.5, 1.5 mM MgCl2, 3mM EDTA, 10% (v/v) glycerol, 0.1% (v/v) NP-40, phosphatase inhibitors (PhosSTOP, Sigma-Aldrich, 04906837001) and protease inhibitor cocktail (Sigma-Aldrich, 05056489001)) with the last wash being on a spin column (Thermo Fisher Scientific, 69705). Immunoprecipitates were eluted using 1 mg/mL 3xFLAG peptide dissolved in IP wash buffer by incubation for 1 h at 4°C. FLAG elution were fractionated on SDS-PAGE and analysed by western blot.

### Detection of ubiquitylated RPB1 using Dsk2

GST-Dsk2 affinity resin was prepared as previously described (Tufegdzic Vidakovic et al., 2019). One Shot BL21 (DE3) Star bacteria transformed with pGEX3-Dsk2 plasmid were grown and 10 ml overnight culture was used to inoculate a 250 ml culture grown to OD_600_ = 0.6, all at 37°C in LB with 100 μg/ml Ampicillin and shaking. Expression was induced with 1 mM IPTG and bacteria grown at 30°C with shaking for 4 hours, cells were then pelleted and snap frozen. The pellet was resuspended in 40ml PBS with protease inhibitors and sonicated (Branson Digital Sonifer 250) at 30% output for 15 s ON/30 s OFF pulses for a total of 10 min ON time. Triton X100 was added to 0.5%, mixed gently and incubated on ice for 30 min. Following a 12000 g, 4°C, 10 min spin the supernatant (lysate) was taken and DTT added to 2 mM final concentration. 1 ml Glutathione Sepharose 4B Beads (GE17-0756-01) were spun at 700 g, washed twice in PBS, added to the lysate and incubated at 4°C with rotation for 4 h. Beads were spun at 700 g, washed for 5 min twice with ice cold PBS + 0.1% Triton X100 then once with ice cold PBS. 1 ml GST-DSK2 beads were resuspended in 3 ml PBS and stored at 4°C before use. For chromatin experiments HEK293 Flp-In T-REx Wild Type and *STK19-*KO clone B3 cells were plated in 2 x 15cm dishes per condition. When around 80% confluent, media was removed and cells were irradiated with 15 J/m^2^ UV and the same media was replaced. For 0 h timepoints media was removed and replaced without irradiation. At the time points indicated cells were scraped from plates and spun down with all media in 50ml Falcon tubes at 300 g, the pellet was washed with PBS and transferred to a 2 ml Eppendorf tube, spun at 300 g and snap frozen before performing chromatin fractionation. A 1% input was taken from the chromatin fraction. 120 μl per sample of bead suspension (30 μl packed bead volume, GST-Dsk2 affinity resin) was spun down at 700 g and resuspended in equivalent volume of buffer. 120 μl was added to each sample and rotated overnight at 4°C. Beads were spun down at 700 g, washed with 5 min, 4°C, rotating incubations twice in TENT buffer (TrisHCL pH7.4 50 mM, EDTA 2 mM, NaCl 150 mM, Triton X100 1%) and once in PBS. 50 μl 1x Sample Buffer was added and the sample boiled for 2 min. 1% input and 20% sample were run on a 4-15% TGX gel and normal western blotting procedure followed.

### Flp-In T-REx HEK293 Cell fractionation

Cell fractionation was carried out as described previously (Gregersen et al., 2019) with some alterations when used for DSK pulldowns - in all buffers 10 % glycerol reduced to 1 %, NP-40 replaced with 1 % Triton X100 and 2 mM NEM added (200 mM stock in ethanol made fresh). All buffers contained cOmplete protease inhibitor (Sigma, 5056489001) and phosSTOP phosphatase inhibitor (Sigma, 4906837001). Pellets were resuspended in 500 μl hypotonic buffer (10 mM HEPES pH 7.5, 10 mM KCl, 1.5 mM MgCl_2_) and incubated on ice for 15 min, homogenised with 20 strokes using a loose pestle and spun at 3000 g, 4°C, 15 min to pellet nuclei. Nuclear pellets were resuspended in 500 μl nucleoplasmic extraction buffer (20 mM HEPES pH 7.9, 1.5 mM MgCl_2_, 150 mM Potassium Acetate, 1 % Glycerol, 1% Triton X100) and incubated on ice for 20 min, then spun at 20 000 g, 4°C, 20 min to pellet chromatin. Chromatin pellets were resuspended in 300 μl chromatin digestion buffer (20 mM HEPES pH 7.9, 1.5 mM MgCl_2_, 150 mM NaCl, 1 % Glycerol, 1% Triton X100) containing Benzonase (MerckMillipore 70746-4)to 75μg/ml (or 1:1000 BaseMuncher (Expedeon, BM0100) for DSK pull-downs) and incubated for 1h, 4°C, rotating, then spun at 20 000 g, 4°C, 20 min, supernatant was kept as the low salt chromatin fraction. Pellets were resuspended in 150 μl high salt chromatin extraction buffer (20 mM HEPES pH 7.9, 500 mM NaCl, 3 mM EDTA, 1.5 mM MgCl_2_, 1 % Glycerol, 1% Triton X100) and incubated on ice for 20 min. 350 μl high salt dilution buffer (20 mM HEPES pH 7.9, 3 mM EDTA, 1.5 mM MgCl_2_, 1 % Glycerol, 1% Triton X100) was added and samples spun at 20 000 g, 4°C, 15 min. Supernatant was pooled with low salt chromatin fraction to form the chromatin fraction. Protein concentration was measured with Bradford assay and each sample adjusted with equivalent buffer (6:3:7 chromatin digestion buffer: high salt chromatin extraction buffer: high salt dilution buffer) to 750 μl at 1 mg/ml.

### RBP1 degradation and stability experiments

To assess the dynamics of total RBP1 and on chromatin after UV irradiation, Flp-In T-REx HEK293 cells were irradiated with 15 J/m^2^ UV-C using an custom built conveyor belt at a confluency of 60% and samples were taken at the indicated times. For RBP1 stability experiments, 100 μM 5,6-dichloro-1-β-D-ribofuranosylbenzimidazole (DRB) was added to cells immediately after UV irradiation, preventing the release of new RNAPIIs into the elongation phase, and allowing us to monitor the fate of RNAPII molecules already engaged in elongation prior to treatment (S2-phosphorylated).

### Recovery of RNA synthesis (RRS)

Cells were irradiated with UV-C light (12 J/m^2^), allowed to recover for the indicated periods, and pulse-labeled with 400 µM 5-ethynyl-uridine (EU; Jena Bioscience) for 1 h followed by a 15 min medium-chase with DMEM without supplements. Cells were fixed with 3.7% formaldehyde in phosphate buffered saline solution (PBS) for 15 min, permeabilized with 0.5% Triton X-100 in PBS for 10 min at room temperature, and blocked in 1.5% bovine serum albumin (BSA, Thermo Fisher) in PBS. Nascent RNA was visualized by click-it chemistry, labeling the cells for 1 h with a mix of 60 µM Atto azide-Alexa594 (Atto Tec), 4 mM copper sulfate (Sigma), 10 mM ascorbic acid (Sigma) and 0.1 μg/mL DAPI in a 50 mM Tris-buffer (pH 8). Cells were washed extensively with PBS and mounted in Polymount (Brunschwig).

### Quantitative PCR (qPCR)

Total RNA was extracted using the RNeasy kit (QIAGEN, 74104) for nascent and mature RNA, following the instructions of the manufacturer including an on-column DNase treatment (QIAGEN, 79254). Reverse transcription was performed using TaqMan Reverse Transcription Reagents (Thermo Fisher Scientific, N8080234). For detection of nascent transcripts, random hexamers were used for the reverse transcription step. cDNA was amplified using iTaq Universal SYBR Green Supermix (BioRad, 172-5124) with 30 cycles of 15 s denaturation at 94°C, 15 s annealing at 58°C, and 20 s extensions at 72°C. Primers amplifying mature GAPDH were used as normalization control. Unless differently stated, ΔCT values were calculated relative to GAPDH before normalizing to the expression level in control sample and experiments were done in triplicate. Primers to amplify nascent RNA were spanning genomic exon-intron regions.

### Unscheduled DNA synthesis

Unscheduled DNA synthesis (UDS) was visualized and quantified using the protocol described before (van den Heuvel et al., 2023; van der Meer et al., 2023). Briefly, cells were plated in DMEM supplemented with 8-10% FCS, followed by serum starvation in DMEM without FCS for at least 24 h to reduce the number of cells going into replication and remove the excess of available deoxy-uridine in the culture medium. Cells were locally UV irradiated through 5 μm pore filters (Milipore; TMTP04700) with 30 J/m^2^ for global genome repair UDS (GGR-UDS) or 100 J/m^2^ for transcription-coupled repair UDS (TCR-UDS), and immediately pulse-labeled with 20 µM 5-ethynyl-deoxy-uridine (EdU; VWR) and 1 µM FuDR (Sigma–Aldrich) for 1 h (GGR-UD) or 4 h (TCR-UDS). After medium-chase with DMEM containing 10 µM Thymidine for 30 min, cells were fixed with 3.7% formaldehyde in PBS for 15 min at room temperature and stored in PBS. Next, cells were permeabilized for 20 min in PBS with 0.5% Triton-X100 and blocked with 3% BSA (Thermo Fisher) in PBS. The incorporated EdU was visualized by click-it chemistry, labeling the cells for 1 h with a mix of 6 µM atto azide-Alexa 647 (Atto Tec), 4 mM copper sulfate (Sigma) and 10 mM ascorbic acid (Sigma) in a 50 mM Tris-buffer (pH 8). After this, the cells were post-fixed with 2% formaldehyde for 10 min and blocked with 100 mM Glycine. Cells were washed extensively with PBS, DNA was denatured with 0.5 M NaOH for 5 min, blocked with 10% BSA (Thermo Fisher) in PBS for 15 min and equilibrated in 0.5% BSA and 0.05% TritonX100 in PBS (WB-buffer). Damaged areas were visualized by labeling the cells for 2 h with mouse anti-CPD (Cosmo Bio in WB-buffer). After primary antibody incubation, cells were washed extensively with WB-buffer, stained with goat anti-rabbit IgG-Alexa 555 (Thermo Fisher) in WB-buffer for 1 h, again washed extensively with WB-buffer, counterstained with 0.1 μg/mL DAPI, washed extensively with PBS and mounted in Polymount (Brunschwig).

### Trabectedin incision assay

Cells were treated with 10 nM trabectedin (MedChemExpress) for 4 h. During the last 15 min, 20 µM 5-Ethynyl-2’-deoxyuridine (5-EdU; Jena Bioscience) was added. Cells were then washed with 300 mM sucrose (Merck) in PBS on ice, pre-extracted with 300 mM sucrose and 0.25% Triton X-100 in PBS for 2 minutes on ice and fixed with 3.7% formaldehyde in PBS for 15 min at room temperature. Cells were subsequently permeabilized with 0.5% Triton X-100 in PBS for 10 min at room temperature, and blocked in 3% bovine serum albumin (BSA, Thermo Fisher) in PBS. Dividing cells were visualized by click-it chemistry, labeling the cells for 30 minutes with a mix of 6 µM Atto azide-Alexa594 or Atto azide-Alexa647 (Atto Tec), 4 mM copper sulfate (Sigma) and 10 mM ascorbic acid (Sigma) in a 50 mM Tris-buffer (pH 8). After washing with PBS, cells were blocked with 100 mM glycine (Sigma) in PBS for 10 min at room temperature and subsequently with 0.5% BSA and 0,05% tween 20 in PBS for 10 minutes at room temperature. To visualize γH2AX, cells were incubated with a primary antibody for phospho-Histone H2A.X Ser139 (JBW301, Merck) for 2 h at room temperature followed by a secondary antibody Anti-Mouse Alexa 555(A-21424, Thermo Fisher) or Anti-Mouse Alexa 647 (A-21235) and DAPI for 1 h at room temperature and mounted in Polymount (Brunschwig).

### DRB run-off microscopy assay

RPE1 cells knockout for XPC, XPC/CSA, or XPC/STK19 stably expressing flag-bio-tagged DDB2 were plated in DMEM supplemented with 8-10% FCS, followed by serum starvation in DMEM without FCS for at least 24 h to reduce the number of replicating cells. Cells were locally UV irradiated through 5 μm pore filters (Milipore; TMTP04700) with 100 J/m^2^ and immediately treated with 100 µM DRB (Sigma, D1916) for the indicated times, washed with PBS and fixed with 3.7% formaldehyde in PBS for 15 min at room temperature. Cells were then permeabilized with 0.5% Triton X-100 in PBS for 10 min at room temperature. After washing with PBS, cells were blocked with 100 mM glycine (Sigma) in PBS for 10 min at room temperature and subsequently with 0.5% BSA and 0,05% tween 20 in PBS for 10 minutes at room temperature. To visualize elongating RNAPII, cells were incubated with a primary antibody for RNAPII-S2 (ab5095, Abcam) for 2 h at room temperature and then with a secondary antibody Anti-Mouse Alexa 555 (A-21424, Thermo Fisher), as well as a Streptavidin/Alexa Fluor™ 488 conjugate recombinant protein (S11223, Thermo Fisher) to detected flag-bio-DDB2 to mark damage sites and DAPI for 1 h at room temperature. Cells were subsequently mounted in Polymount (Brunschwig).

### Microscopic analysis of fixed cells

Images of fixed samples were acquired on a Zeiss AxioImager M2 or D2 widefield fluorescence microscope equipped with 63x PLAN APO (1.4 NA) oil-immersion objectives (Zeiss) and an HXP 120 metal-halide lamp used for excitation. Fluorescent probes were detected using the following filters for DAPI (excitation filter: 350/50 nm, dichroic mirror: 400 nm, emission filter: 460/50 nm), GFP/Alexa 488 (excitation filter: 470/40 nm, dichroic mirror: 495 nm, emission filter: 525/50 nm), Alexa 555/594 (excitation filter: 545/25 nm, dichroic mirror: 565 nm, emission filter: 605/70 nm), or Alexa 647 (excitation filter: 640/30 nm, dichroic mirror: 660 nm, emission filter: 690/50 nm). Images were recorded using ZEN 2012 software (blue edition, version 1.1.0.0) and analysed in Image J (1.48v).

### Trabectedin-COMET chip assay

Cells were enriched at the G1 phase by incubating in growth medium supplemented with 2 μM palbociclib (hereafter referred to as the ‘working medium’) for 24 h prior to exposure to trabectedin. Following this, cells were embedded in a 30 μm COMET chip (catalog no. 4250-096-01, Trevigen) and further incubated for 30 min at 37 °C in the working medium. The medium was removed, and cells were incubated in the working medium supplemented with 50 nM trabectedin for 2 h. Following trabectedin wash-out, cells were incubated in the working medium for 0, 2 or 4 h. A 30 μm COMET chip (catalog no. 4250-096-01, Trevigen) was equilibrated in 100 mL tissue culture grade 1X PBS for 30 min at room temperature and placed into the 96-well COMET chip System (catalog no. 4260-096-CS, Trevigen). A single cell suspension was prepared in 6 mL working medium at 1.0×10^5^ cells/mL. Single cell suspension was aliquoted 100 µL per well. The lid-covered COMET chip System was placed in the tissue culture incubator for 10 min, gently rocked E-W and N-S, and this was repeated 3 times. The working medium was aspirated carefully not to remove cells. Cells were treated with trabectedin as described above. The COMET chip with treated cells was then overlaid with 6 mL of 1% low melting agarose (catalog no. 4250-500-02, Trevigen), followed by cell lysis with 50 mL lysis solution (catalog no. 4250-500-01, Trevigen) for overnight at 4 °C. Unwinding of DNA was carried out for 30 min twice in 250 mL alkaline solution (200 mM NaOH, 1 mM EDTA, 0.1% Triton X-100). Electrophoresis was carried out for 50 min, 1 V/cm at 4 °C in 700 mL alkaline solution. After electrophoresis, the COMET chip was neutralized for 15 min twice at 4 °C in 100 mL 0.4 M Tris pH 7.4 and equilibrated for 30 min at 4 °C in 100 mL 20 mM Tris pH 7.4, followed by staining in 50 mL 0.2 X SYBR Gold (catalog no. S11494, Invitrogen) at room temperature for 2 h. The stained COMET chip was destained at room temperature in 100 mL 20 mM Tris pH 7.4 up to 1 h. Comets were imaged with 4X magnification on a fluorescence microscope (BX53, Olympus). % DNA in tail was quantified with Comet analysis software (catalog no. 4260-000-CS, Trevigen) (Son et al., 2024).

### Protein expression and purification

STK19 DNA fragment was produced by IDT after sequence optimization for expression in *E. coli.* DNA fragment was cloned into the 1C cloning vector (addgene no. 29654) by ligation-independent cloning (Gradia et al., 2017).Components of the RNAPII-TCR complex were cloned, expressed and purified as previously described (Bernecky et al., 2016; Kokic et al., 2021; Kokic et al., 2024). Human STK19 was expressed in *E. coli* BL21 DE3 cells in LB media. Cells were grown at 37 °C until they reached the OD600 of 0.8, followed by induction with 1 mM IPTG and protein expression overnight at 18 °C. Cells were harvested by centrifugation at 6000 rpm for 20 min and resuspended in lysis buffer (400 mM NaCl, 20 mM Tris-Cl pH 7.9, 10% glycerol, 1mM DTT, 30 mM imidazole, 0.284 µg/mL leupeptin, 1.37 µg/mL pepstatin A, 0.17 mg/mL PMSF, and 0.33 mg/mL benzamidine). Cells were opened by sonication and the lysate was clarified by centrifugation at 20 000 rpm for 1h. Clarified lysate was loaded onto 5 mL HisTrap column (Cytiva) equilibrated in lysis buffer. Column was washed with 10 CV lysis buffer and the protein was eluted with a 0-100% elution buffer (as lysis buffer with 500 mM imidazole) over 5CV. Peak protein fractions were supplemented with TEV protease and dialyzed overnight against the dialysis buffer (400 mM NaCl, 20 mM HEPES pH 7.5, 10% glycerol and 1mM DTT). Digested protein was passed over a 5ml HisTrap column equilibrated in dialysis buffer. The flow-through was collected, concentrated and polished over a Superdex200 10/300 GL size exclusion column (GE Healthcare) pre-equilibrated in dialysis buffer. Peak protein fractions were pooled and concentrated, and protein aliquots flash frozen in liquid nitrogen and stored at -80 °C.

### Electron microscopy

Sample preparation. DNA and RNA sequences used for the preparation of RNAPII elongation complex are: GTA TTC GCT CTG CTC CTT CTCCCA TCC TCT CGA TGG CTA TGA GAT CAA CTA G (template strand), CTA GTT GAT CTC ATA TTT CAT TCC TAC TCA GGA GAA GGA GCA GAG CGA ATA C (non-template strand) and rC rArArA rArUrC rGrArG rArGrG rA (RNA). The RNA and the template strand were annealed in water by heating the solution to 95 °C followed by slow cooling to 4 °C. RNAPII and hybridized DNA:RNA were mixed in 1:1 molar ration, followed by the addition of 1.5x molar access of non-template strand. The solution was next supplemented with CSB, CSA-DDB1, UVSSA, ELOF1 and STK19 in 1:1:1:2:3 molar ratio to RNAPII and the sample was incubated on ice for 1 h. Final complex formation buffer conditions were 100 mM NaCl, 5% glycerol, 2 mM HEPES pH 7.5, 2 mM MgCl_2_ and 1 mM DTT. The RNAPII-TCR-STK19 complex was purified via Superose 6 Increase 3.2/300 column equilibrated in the final complex formation buffer. Peak fractions were pooled and crosslinked with 0.1% glutaraldehyde on ice for 1 min. The reaction was quenched using lysine (50 mM final) and aspartate (20 mM final). The quenched solution was dialyzed in Slide-A-Lyzer MINI Dialysis Device of 20 MWCO (Thermo Fischer Scientific) for 7 h against the complex formation buffer without glycerol. After the dialysis, the protein solution was immediately used for the preparation of cryo-EM grids. 4 µl of the sample was applied onto freshly glow-discharged R2/1 carbon grids (Quantifoil), followed by grid blotting for 4 s and plunging into liquid ethane using a Vitrobot Mark IV (FEI) operating at 4 °C and 100 % humidity. Imaging, data processing and model building. Images of the sample were taken on a 200 keV Thermo Scientific Glacious Cryo-Transmission Electron Microscope using a Falcon 3 direct electron detector. Images were taken at a magnification of 120 000x (1.23 Å per pixel) with a dose of 1.32 e/Å2 per frame over 30 frames by using EPU software. The total of 2696 micrographs were pre-processed in Warp (Tegunov and Cramer, 2019), followed by automated particle picking in the same software. 778 361 particles were imported in CryoSPARC (Punjani et al., 2017) and used for An Initio reconstruction with two classes. The two classes were used as input for heterogeneous refinement. Subsequent rounds of heterogeneous refinement were used to obtain a particle set with stably bound STK19, as outlined in Supplemental Figure 5. Final subset of 113 958 particles were used for NU refinement in CryoSPARC, which resulted in the final map at the overall resolution of 3.3 Å. RNAPII-TCR-ELOF structure (PDB 8B3F) together with the AlphaFold (Jumper et al., 2021) model for human STK19 were rigid-body docked into the refined density in Chimera (Goddard et al., 2018; Pettersen et al., 2004), followed by real-space refinement in PHENIX (Adams et al., 2010) and manual adjustment in Coot (Emsley et al., 2010).

### In vitro XPD helicase assay

Stimulation of the the 5’->3’ helicase activity of TFIIH by XPA and STK19 was monitoring as described previously (Kokic et al., 2019). The labeled 5′-FAM DNA oligonucleotide (FAM-5′-TTCACCAGTGAGACGGGCAACAGC-3′) was annealed with a 1.5-fold excess of the quenching DNA oligonucleotides (5’-AAATCGCGCGTCTAGACTCAGCTCTGGCTGTTGCCCGTCTCACTGGTGAA-3’-BHQ) in 10 mM Tris-HCl (pH 7.5) and 50 mM NaCl. For each reaction, 0.4 pmol of DNA duplex, 8 pmol of core TFIIH (Kim et al., 2022), 100 mM KCl, 20 mM HEPES-KOH pH 7.0, 5% glycerol, 0.2 mg/ml bovine serum albumin, 3 mM phosphoenolpyruvate, 10 mM MgCl_2_, 1 mM DTT, and 2.5 unit pyruvate kinase (Sigma) in a total volume of 20 μl were pre-incubated at 26°C for 10 min. ATP (2 mM final) was added to each reaction, and the unwinding was monitored at 26°C by using the Synergy NEO2 Hybrid Multi-Mode Reader (Biotek) with an excitation wavelength of 495 nm and an emission wavelength of 520 nm. The percentage of unwound product was calculated by dividing the observed total fluorescence intensity by the intensity of the fluorescent oligo in the reaction buffer (mimicking fully unwound DNA). To measure the stimulation of unwinding of TFIIH by STK19 and XPA, the reaction was supplemented with STK19 (1.2 µM or 2.4 µM) or XPA (1.2 µM).

### CRISPR screen

RPE1-iCas9 cells (van der Weegen et al., 2021) were transduced at an M.O.I. of ∼0.2 with a 1:1000 dilution of TKOv3-in-pLCKO lentiviral library in medium containing 8 µg/mL hexadimethrine bromide (Sigma-Aldrich). The library was a gift from Katherine Chan, Amy Tong and Jason Moffat (Donnelly Centre, University of Toronto, Toronto, ON). 24 h after transduction, puromycin (Sigma-Aldrich) was added to 5 μg/mL to select for transduced cells. After all cells in non-transduced control populations had died and dishes with transduced populations had reached 90% confluence, a t=0 sample was taken for each of the three populations. From the remaining cells of each population 30×10^6^ (corresponding to a library representation of >400) were grown as a control population, and additionally 30×10^6^ were grown in the presence of 4-NQO at a concentration of 100 ng/ml. Doxycycline was added to the medium of all replicates from t=0 onwards to induce expression of Cas9, at a concentration of 200 ng/mL. After three doublings 30×10^6^ cells of each population were passed. After 12 doublings, all populations were harvested.

### Sequencing and analysis of the CRISPR screen

Genomic DNA was isolated from each population using the Blood and Cell Culture DNA Maxi Kit (Qiagen). 3 µg gDNA of each population was amplified using the KAPA HiFi ReadyMix PCR Kit (Roche) with the TKO outer Fw and Rv primers (Primers are listed in Supplementary Table 4) followed by a second PCR reaction using reverse primers with different Illumina i7 index sequences for each sample to identify the sample after pooled sequencing as described43. The second PCR products of each pool were purified by QIAquick PCR Purification Kit (Qiagen). RNAseq samples were added to 10% to create balanced reads for the first 21 nucleotides sequenced. Up to twelve samples were sequenced in a single HiSeq4000 lane in an SR50 run using standard reagents and conditions and reads were mapped to the TKOv3 library sequences, not allowing any mismatches. The screen data was analyzed using DrugZ version 1.1.0.233, Log2_Fold-Change was calculated by first normalizing all samples to an equal number of total reads (pseudocount +1), then calculating the median fold change per guide for the three replicate screen samples, and finally calculating the Log2 of the median fold change of all guides targeting the gene. For network analysis, the physical interactions of differentially depleted genes were extracted from the gene interaction database, GeneMANIA. The string categories TC-NER, NER, and BER are combined as the DNA Repair category, while the rest are categorized as having an unknown protective function. The sizes of the nodes represent the most significant -Log false discovery rate (FDR), and the inner color of the node represents the most significant fold change (F2C) among the 4NQO and IlludinS screens. The color of the outer layers of the nodes represents the screen in which they were identified.

### TCR-seq

Cells were grown to 80-90% confluency and crosslinked with 0.5 mg/mL disuccinimidyl glutarate (DSG; Thermo Fisher) in PBS for 45 min at room temperature. Cells were washed once with PBS followed by incubation with 1% PFA for 20 min at RT. Fixation was stopped by addition of glycine in PBS to a final concentration of 0.1 M for 3 min at RT. This was followed by washing with cold PBS and collection of the cells in 0.25% Triton X-100, 10 mM EDTA (pH 8.0), 0.5 mM EGTA (pH 8.0), and 20 mM Hepes (pH 7.6) in miliQ. Chromatin was pelleted by centrifugation for 5 min at 400 × g and incubated in 150 mM NaCl, 1 mM EDTA (pH 8.0), 0.5 mM EGTA (pH 8.0) and 50 mM Hepes (pH 7.6) in miliQ for 10 min at 4 °C. Chromatin was again pelleted by centrifugation and resuspended in ChIP-buffer (0.15 % SDS, 1 % Triton X-100, 150 mM NaCl, 1 mM EDTA (pH 8.0), 0.5 mM EGTA (pH 8.0) and 20 mM Hepes (pH 7.6) in miliQ) to a final concentration of 15 × 10^6^ cells/mL. Chromatin was sonicated to approximately one nucleosome using the Bioruptor Pico (Diagenode), 8 cycles of 30 s ON/30 s OFF in a 4°C water bath. ChIP was performed on 280-300ul sonicated material (∼25 µg of chromatin per sample) with 3 µg of rat anti-RNAPII-S2 antibody (Millipore, clone 3E10) by overnight incubation at 4 °C, followed by a 1.5 h protein-chromatin pull-down with 20 µl protein G Dynabeads (Thermo Fisher, 10003D). ChIP samples were washed extensively, followed by decrosslinking for 4 h at 65 °C in the presence of proteinase K. DNA was purified using a Qiagen MinElute kit. Sample libraries were prepared using Hifi Kapa sample prep kit and A-T mediated ligation of Nextflex adapters or xGen UDI-UMI adapters. Samples were sequenced using an Illumina NextSeq500 or HiSeq X, using paired-end sequencing with 42 or 151 bp from each end. After sequencing, a quality profile was generated for each sample, using FastQC (Version 0.11.9). Sequences were trimmed using TrimGalore (Version 0.6.5). Reads were aligned to the Human Genome 38 using bwa-mem tools (BWA (Version 0.7.17) and reference genome GCA_000001405.15_GRCh3847. Only uniquely mapping and high-quality reads (> q30) were included in the analyses. Duplicate reads were removed using Samtools (Version 1.11) with fixmate -m and markdup -r settings. Bam files were converted into stranded TagDirectories and UCSC genome tracks using HOMER tools (Version 4.8.2) (Heinz et al., 2010). Example genome tracks were generated in IGV (Version 2.4.3). Stranded metagene profiles were generated for the 3,000 actively transcribed genes defined in our BrU-seq data, using the makeMetaGeneProfile.pl tool of HOMER with default settings.

### BrU-seq

RPE1 cells were plated in 3x 15-cm plates per condition and grown to 80-100% confluency. Subsequently, cells were either mock-treated or irradiated with UV-C light (12 J/m^2^). Cells were then incubated in conditioned media for different periods of time (0, 3 and 16 h) before incubation with 2 mM bromouridine (BrU, Sigma) at 37 °C for a 30 min. The cells were then lysed in TRIzol reagent (Invitrogen) and BrU-containing RNA was isolated as previously described (Andrade-Lima et al., 2015; van den Heuvel et al., 2021a; van der Weegen et al., 2021). cDNA libraries were made from the BrU-labeled RNA using the Illumina TruSeq library kit and paired-end 151 bp sequenced using the Illumina NovaSeq platform at the University of Michigan DNA Sequencing Core. For each BrU-seq sample, a sequencing quality profile was generated using FastQC (Version 0.11.9). Sequences were trimmed using TrimGalore (Version 0.6.5). Reads were aligned to the Human Genome 38 using STAR (Version 2.7.7a) with genome file GCA_000001405.15_GRCh38 and splice junction file Homo_sapiens.GRCh38.94.correctedContigs.gtf. Bam files were converted into stranded TagDirectories and UCSC genome tracks using HOMER tools (Version 4.8.2) (Heinz et al., 2010). Example genome tracks were generated in IGV (Version 2.4.3). A list of 49,948 gene-coordinates was obtained from the UCSC genome database (https://genome.ucsc.edu/cgi-bin/hgTables) selecting the known Canonical table containing the canonical transcription start sites per gene. To prevent contamination of binding profiles, genes were selected to be non-overlapping with at least 2 kb between genes and a minimal size of 3 kb (n=9,944). From this, a set of 3,000 actively transcribed genes was selected based on the BrU-seq signal in the first 3 kb of genes in WT cells, using the AnnotatePeaks.pl tool of HOMER with default settings. These 3,000 actively transcribed genes were used in downstream analyses, unless stated otherwise. Read-densities around transcription start sites (TSSs) were quantified using the AnnotatePeaks.pl tool of HOMER using default settings and 200bp resolution. Individual datasets were subsequently processed into heatmaps or density profiles using R (Version 4.0.5) and Rstudio (Version 1.1.423) (Team, 2019). To quantitatively compare different read-density profiles, each profile was normalized to nascent transcript levels, as quantified by 5-EU labeling (RRS protocol), relative to the control in their specific cell type. For example, RPE1 iCas9 cells 3 h after UV irradiation showed 32% nascent transcription relative to mock-treated RPE1 iCas9 cells, so we multiplied the read-density in each 200-bp bin by 0.32.

### TT_chem_-seq

TT-seq was performed essentially as described (Gregersen et al., 2020; Gregersen et al., 2019) Biological triplicates were generated for each condition. Flp-In T-REx HEK293, wilt type or *STK19-*KO cells were irradiated with 15 J/m2 UV-C and in vivo labeling of nascent RNA was achieved with 1 mM 4SU (Glentham Life Sciences, GN6085) pulse for exactly 15 min at the appropriate time to fit with the specified time points. Labeling was stopped by TRIzol (Thermo Fisher Scientific, 15596026) and RNA extracted accordingly to the manufacturer’s instructions. As a control for equal sample preparation, S. cerevisiae (strain BY4741, MATa, his3D1, leu2D0, met15D0, ura3D0) 4-thiouracil (4TU)-labeled RNA was spiked in to each sample. S. cerevisiae were grown in YPD medium overnight, diluted to an OD600 of 0.1, and grown to mid-log phase (OD600 of 0.8) and labeled with 5 mM 4TU (Sigma-Aldrich, 440736) for 6 min. Total yeast RNA was extracted using the PureLink RNA Mini kit (Thermo Fisher Scientific, 12183020) following the enzymatic protocol. For purification of 4SU labeled RNA, 100 μg human 4SU labeled RNA was spiked-in with 1/100 of 4TU-labeled S. cerevisiae RNA. The 101 μg RNA (in a total volume of 100 uL) was fragmented by addition of 20 μL 1 M NaOH and left on ice for 20 min. Fragmentation was stopped by addition of 80 μL 1 M Tris pH 6.8 and cleaned up twice with Micro Bio-Spin P-30 Gel Columns (BioRad, 7326223) according to the manufacturer’s instructions. Biotinylation of 4SU- and 4TU-residues was carried out in a total volume of 250 μl, containing 10 mM Tris-HCl pH 7.4, 1 mM EDTA and 5 mg MTSEA biotin-XX linker (Biotium, BT90066) for 30 min at room temperature in the dark. RNA was then purified by phenol:chloroform extraction, denatured by 10 min incubation at 65°C and added to 200 μL μMACS Streptavidin MicroBeads (Milentyl, 130-074-101). RNA was incubated with beads for 15 min at room temperature and beads applied to a μColumn in the magnetic field of a μMACS magnetic separator. Beads were washed twice with pull-out wash buffer (100 mM Tris-HCl, pH 7.4, 10 mM EDTA, 1 M NaCl and 0.1% Tween20). Biotinylated RNA was eluted twice by addition of 100 mM DTT and cleaned up using the RNeasy MinElute kit (QIAGEN, 74204) using 1050 μL 100% ethanol per 200 μL reaction after addition of 750 μL RLT buffer to precipitate RNA < 200 nt. Libraries for RNA sequencing were prepared using the KAPA RNA HyperPrep Kit (KR1350) with modifications. 75 ng of RNA per sample were mixed with FPE Buffer, but fragmentation procedure was omitted and RNA was instead denatured at 65°C for 5 min. The rest of the procedure was performed as recommended by the manufacturer, with the exception of SPRI bead purifications: after adaptor ligation, 0.95x and 1x SPRI bead-to-sample volume ratios were used (instead of two rounds of SPRI purification with 0.63x volume ratios). This was done to retain smaller (150-300 bp) cDNA fragments in the library which would otherwise be lost in purification. Libraries were amplified using 6 cycles of PCR. The libraries were then sequenced with single end 75 bp reads on the HiSeq 2500, with 50,000 reads per sample.

### TT_chem_-seq data analyses

TT_chem_-seq data were processed using previously published protocol (Gregersen et al., 2020). Reads were trimmed to remove Illumina adapter sequences using cutadapt v1.18 (10.14806/ej.17.1.200) before separate alignment to the Homo sapiens GRCh38 (target) and S. cerevisiae sacCer3 (spike-in) genomes using the STAR v2.5.2a RNA-seq aligner (Dobin et al., 2013) with the “-quantMode GeneCounts” option to quantify gene level abundance against Ensembl release 86 transcript annotations. The yeast gene-level counts were used to calculate per-sample scale-factors necessary for the downstream normalisation of the human data. Counts were imported into R and passed to the estimateSizeFactors function of the Bioconductor package DESeq2 (Love et al., 2014). Human genome normalised coverage was calculated for each sample in a strand-specific manner using BEDTools v2.27.1 genomecov function, providing the reciprocal of the yeast derived scale factor via the “-scale” option, (Quinlan and Hall, 2010). The resulting coverage matrix was then converted to bigwig format at single-bp resolution. Metagene profiles of protein-coding genes from standard chromosomes were created from the normalised bigwig files using Deeptools v2.5.3’s computeMatrixOperations function (Ramirez et al., 2014), stratifying based on gene width (50-150kb, >150kb).

---

**Figure S1.**
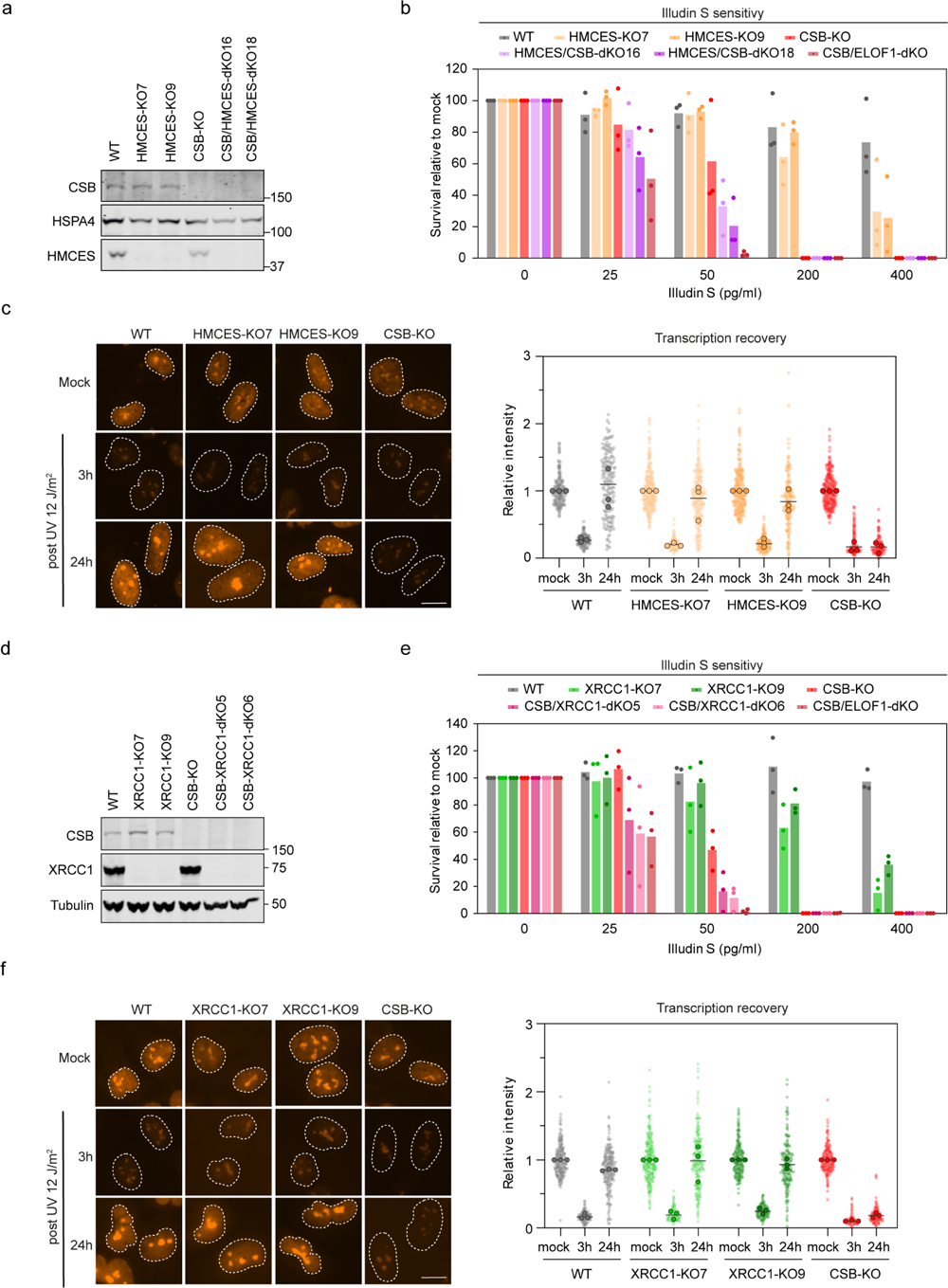
HMCES and XRCC1 confer Illudin S sensitivity but are not TCR factors. **a.** Western blot analysis of parental, single or double *HMCES* knockout RPE1 cells with the indicated genotype. **b.** Quantification of clonogenic survival after Illudin S treatment in *HMCES*-KO RPE1 cells with the indicated genotype. Bars represent the means of 3 experiments, with individual data points shown as circles. **c.** Representative microscopy images (left) and quantification (right) of RPE1 cells with the indicated genotype after 1-h EU labeling. Cells were untreated or analyzed at 3 h or 24 h after irradiation with 12 J/m^2^ UV-C. All cells are depicted as individual semi-transparent data points with the bar representing the mean of all data points. The individual means of three biological replicates are depicted as solid circles with black lines. Scale bar, 10 μm. **d.** Western blot analysis of parental, single or double *XRCC1* knockout RPE1 cells with the indicated genotype. **e.** Quantification of clonogenic survival after Illudin S treatment in *XRCC1*-KO RPE1 cells with the indicated genotype. Bars represent the means of 3 experiments, with individual data points shown as circles. **f.** Representative microscopy images (left) and quantification (right) of RPE1 cells with the indicated genotype after 1-h EU labeling. Cells were untreated or analyzed at 3 h or 24 h after irradiation with 12 J/m^2^ UV-C. All cells are depicted as individual semi-transparent data points with the bar representing the mean of all data points. The individual means of three biological replicates are depicted as solid circles with black lines. Scale bar, 10 μm.

**Figure S2.**
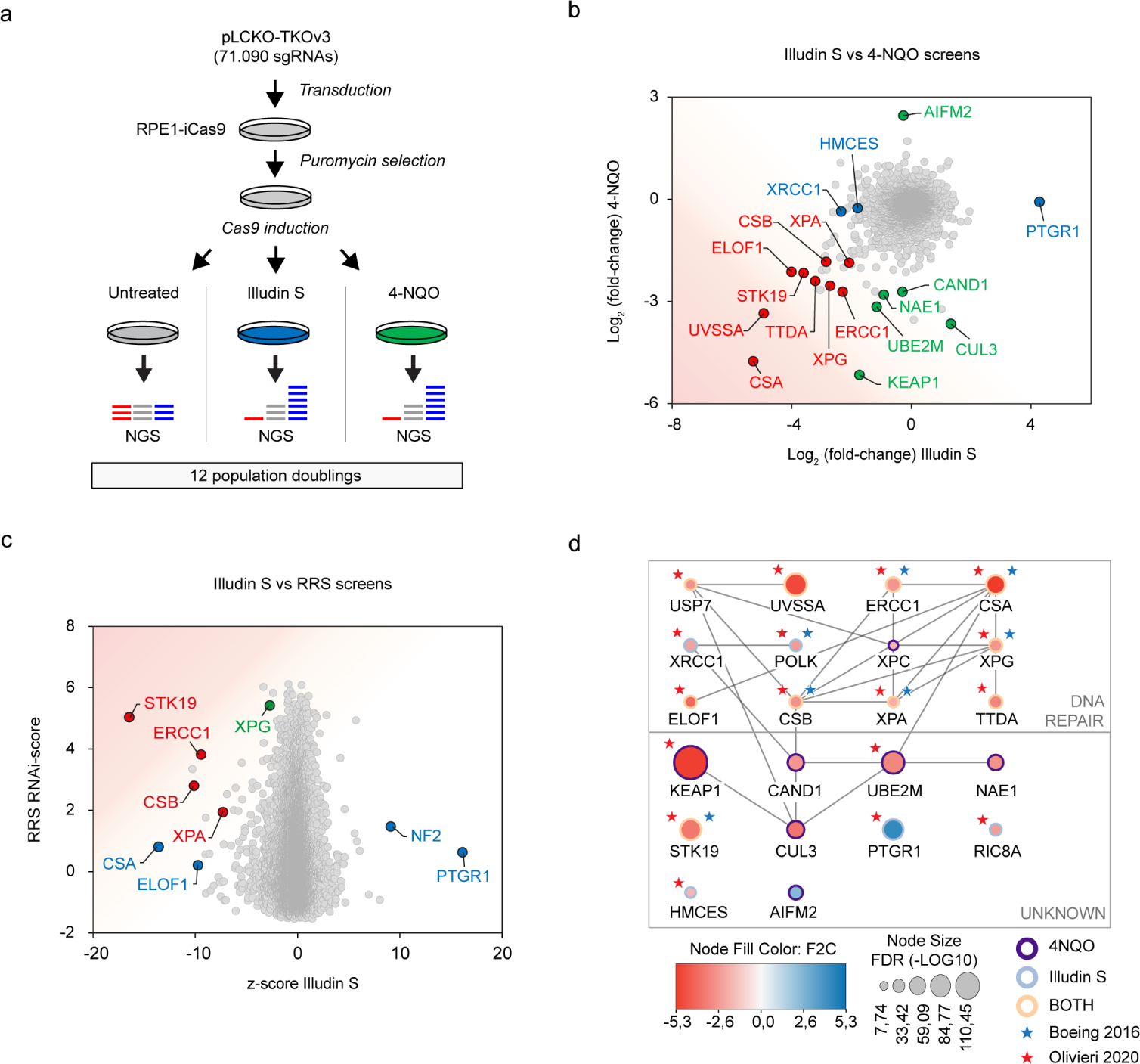
CRISPR screens identify STK19 as a new TCR factor. **a.** Schematic representation of the CRISPR/Cas9 screens in RPE1-iCas9 cells. b. Correlation of the CRISPR screen results showing the fold increase (Log_2_) on Illudin S (x-axis) against the fold increase (Log_2_) on 4-NQO (y-axis). Hits specific for Illudin S or 4-NQO are marked in blue and green, respectively. Hits that scored in both screens are marked in red. c. Correlation of the Illudin S screen z-scores against our previous RRS RNAi-score (Boeing et al., 2016). Hits specific for the Illudin S or RRS screen are marked in blue and green, respectively. Hits that scored in both screens are marked in red. d. Network analysis of highly significant hits representing genes essential for Illudin S or 4-NQO resistance. Grey lines reflect known protein-protein interactions based on Cytoscape and BioGRID. Shared hits with the RRS RNAi screen (Boeing et al., 2016) or a previous Illudin S screen (Olivieri et al., 2020) are marked with a blue or red asterisk, respectively.

**Figure S3.**
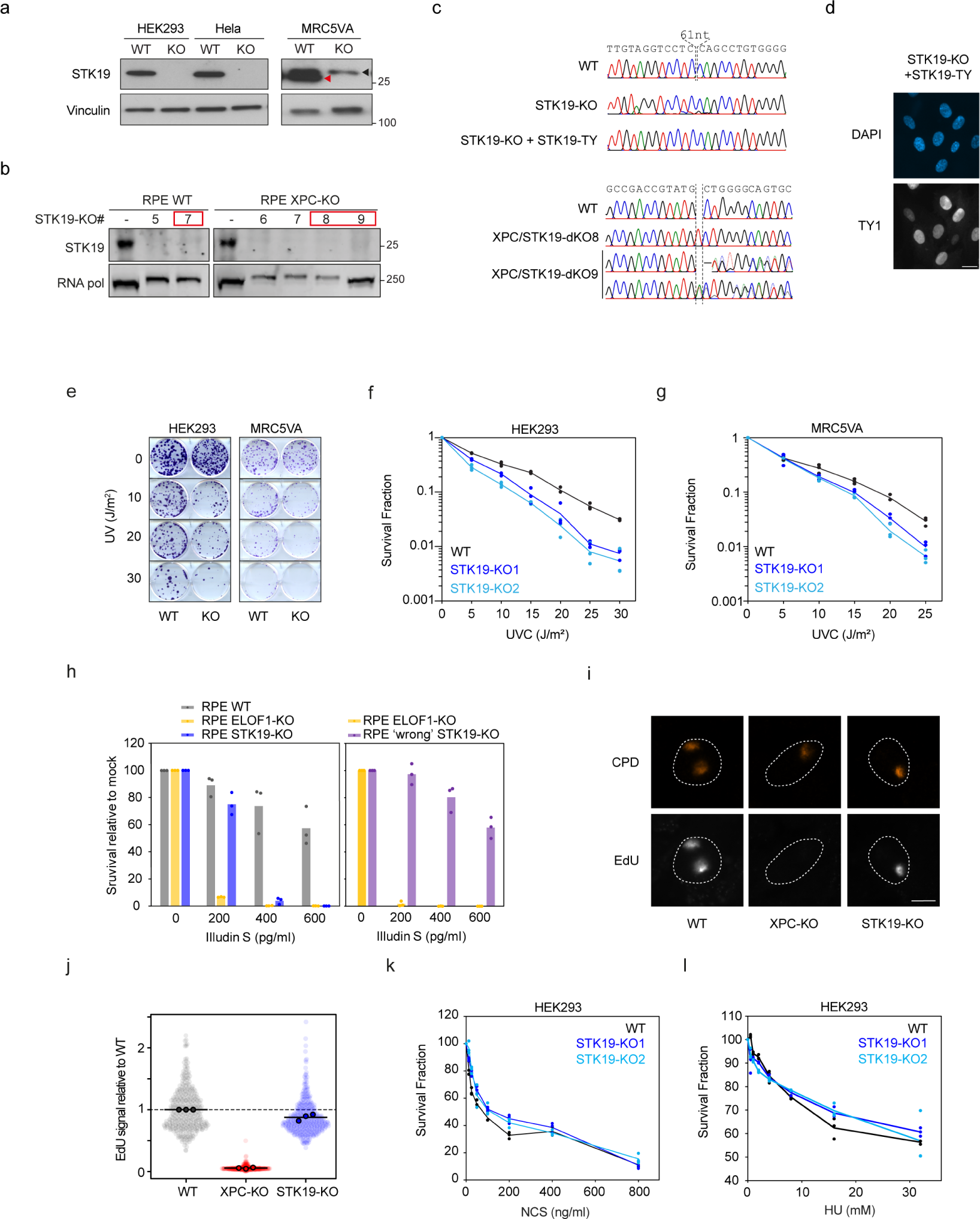
STK19 knockout validation. **a.** Western blots for the validation of *STK19-*KO in Flp-In T-REx HEK293, Flp-In T-REx HeLa and MRC5VA. The red arrow indicates the correct size of STK19, and the black arrow indicates an unspesific band. Vinculin is used as loading control. **b.** Western blot analysis of WT and *STK19-*KO in parental RPE1 or RPE1 *XPC-*KO cells. Selected clones are marked in red. **c.** Sanger sequencing of RPE1 cells (parental or *XPC-*KO) to verify the knockout of STK19. **d.** Nuclear localization of TY1-tagged STK19 protein expressed in *STK19-*KO cells. Scale bar, 20 μm. **e.** Example of the clonogenic assay performed in Flp-In T-REx HEK293 and MRC5VA at different UV doses. **f.** Quantification of the experiments exemplified in **e**. of Flp-In T-REx HEK293. Lines represent the means of 3 experiments with individual data points shown as circles. **g.** Quantification of the experiments exemplified in **e**. of MRC5VA. Lines represent the means of 3 experiments with individual data points shown as circles. **h.** Quantification of clonogenic survival after Illudin S treatment in RPE1 cells with the indicated genotype. The ‘wrong’ *STK19-*KO clone has been transfected with an sgRNA targeting the misannotated exon 1. Bars represent the means of 3 experiments with individual data points shown as circles **i.** Representative images of EdU incorporation in RPE1 cells with the indicated genotype to measure UDS after 100 J/m^2^ UV-C irradiation through 5 µm pore membranes. Sites of local damage are identified by CPD staining. UDS measured in this way mainly reflects GGR activity. Scale bar, 10 μm. **j.** Quantification of UDS signal from **i**. All cells are depicted as individual semi-transparent data points with the bar representing the mean of all data points. The individual means of three biological replicates are depicted as solid circles with black lines. **k.** Quantification of cell survival after treatments with different doses of neocarzynostatin (NCS) in Flp-In T-REx HEK293 (HEK293) WT cells or two *STK19-*KO clones. Cell growth was quantified using luminescence CellTiter-Glo assay. Lines represent the means of 3 experiments with individual data points shown as circles. **l.** Quantification cell survival after treatments with different doses of hydroxyurea (HU) in Flp-In T-REx HEK293 (HEK293) WT cells or two *STK19-*KO clones. Cell growth was quantified using luminescence CellTiter-Glo assay. Lines represent the means of 3 experiments with individual data points shown as circles.

**Figure S4.**
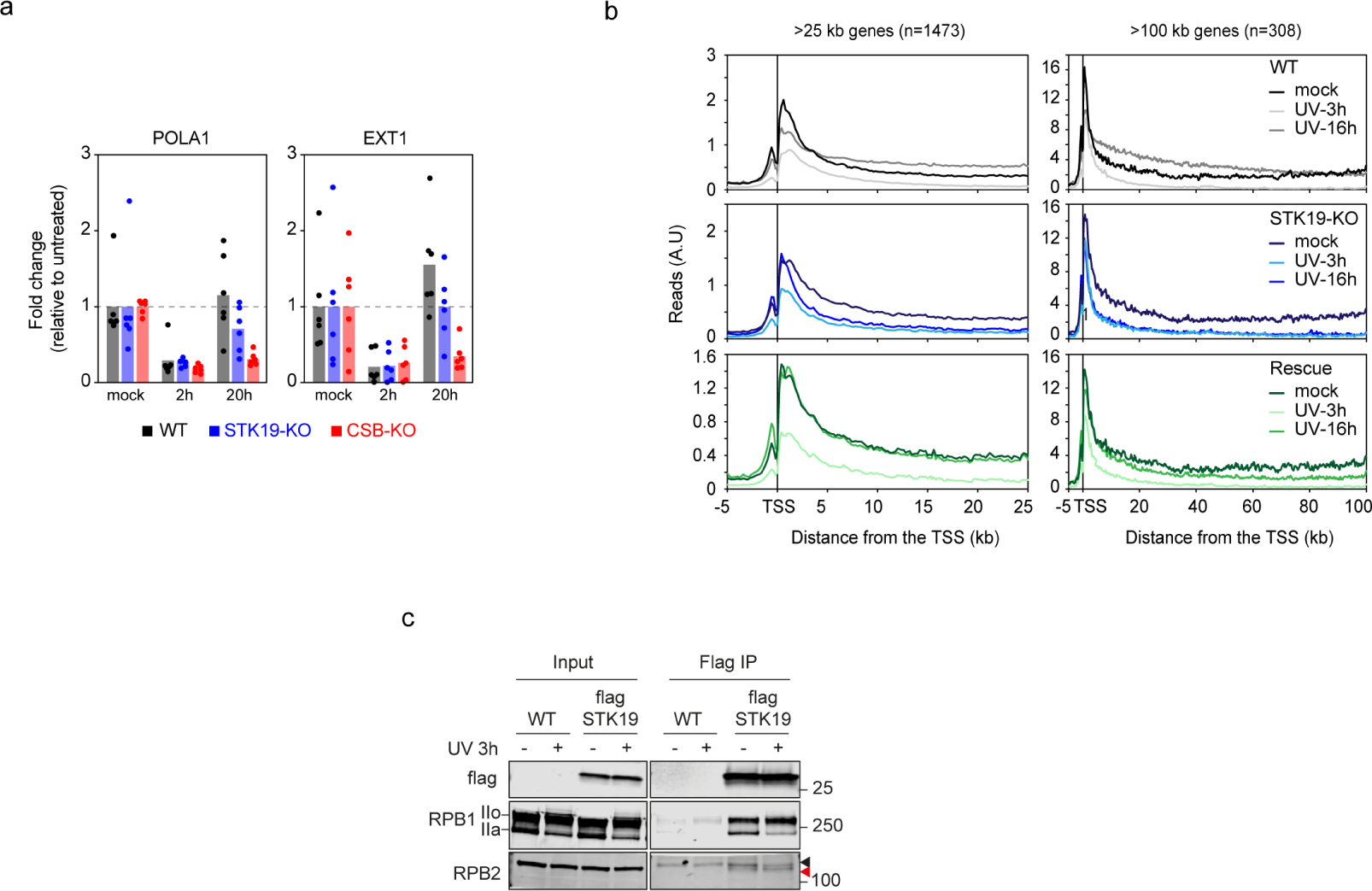
STK19 regulates transcription recovery after UV. **a.** qPCR quantification of *EXT1* and *POLA1* transcription. Primers were located in promoter-proximal regions of indicated genes. WT, *SKT19*-KO and *CSB*-KO knockout cells were either untreated, or collected 2 h and 20h post UV irradiation (15 J/m^2^). Data are normalized to the mature *GAPDH* expression, and to untreated condition for each cell line. Bars represent averages of six biological replicates, with individual datapoints shown as circles. **b.** Average read count profiles from -5 kb to +50 kb around the transcription start site (TSS) of 1.473 genes above 25 kb, and 308 genes above 100 kb in RPE1 cells with the indicated genotype that were untreated or at 3 or 16 h after UV-C irradiation with 12 J/m^2^. **c.** Western blot of the input and immunoprecipitation of FLAG-STK19 before and 3h after UV irradiation. RPB1 and RPB2 are shown as well as Flag for experimental control of STK19 detection. Red arrow indicates the correct band for RPB2 and black arrow a contaminant band detected by the antibody.

**Figure S5.**
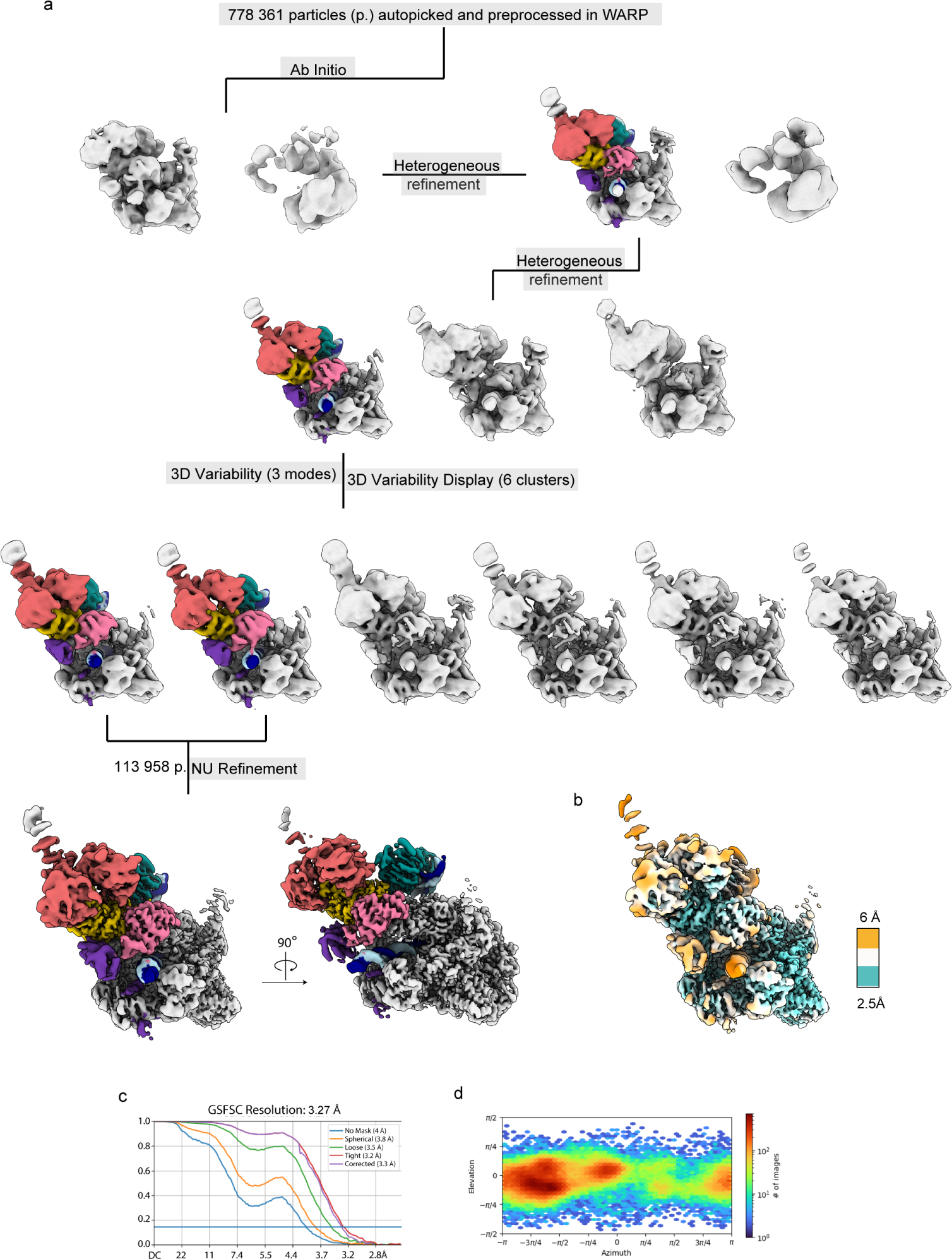
Cryo-EM analysis of the RNAPII-TCR-STK19 complex. **a.** Processing tree. Classes used for further processing are color-coded according to complex subunits, as in Figure 4c. **b.** Local resolution estimate for the final map. **c.** Fourier shell correlation plots for the final map. **d.** Angular distribution plot for particles contributing to the final map.

**Figure S6.**
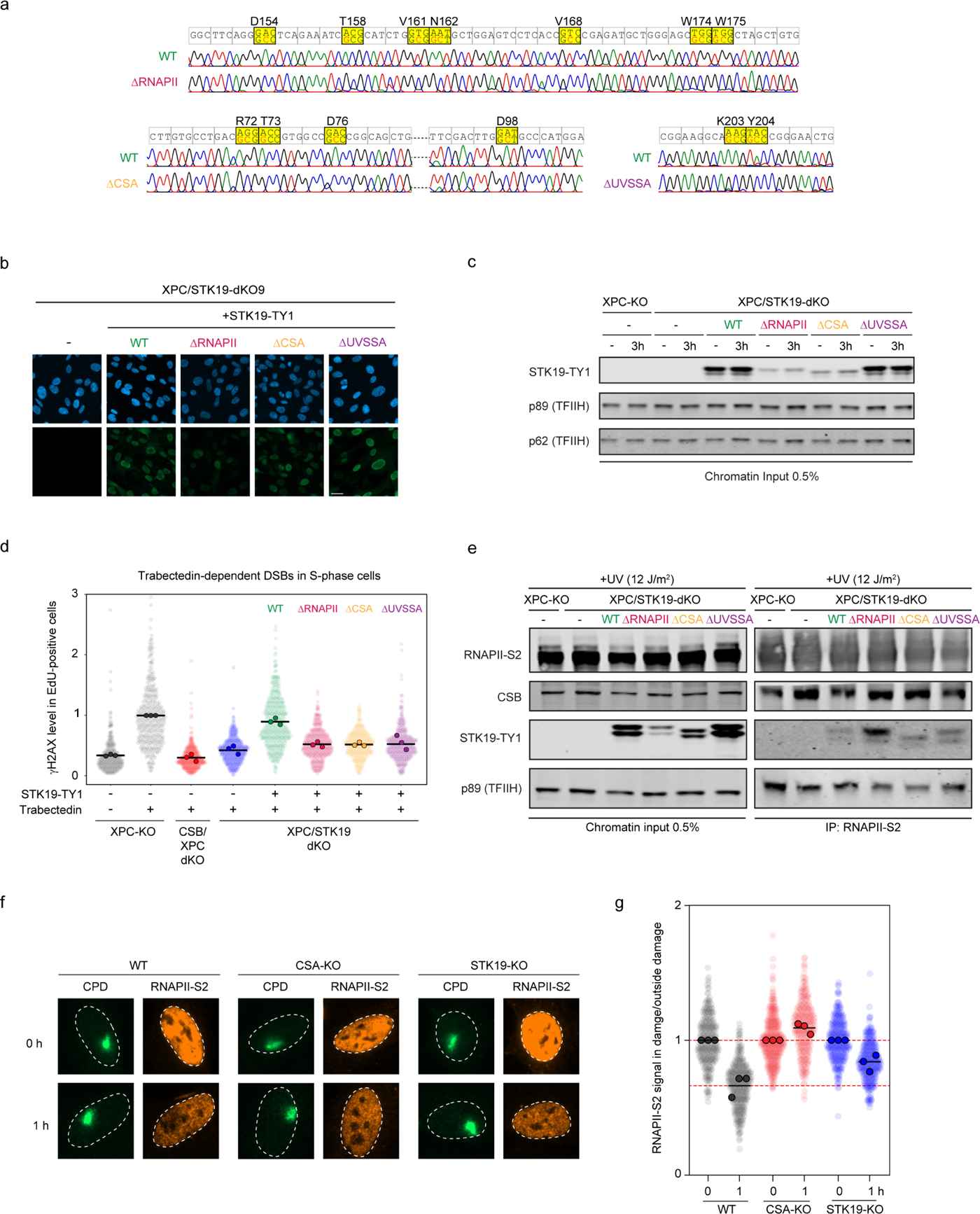
Validation of STK19 mutants. **a.** Sanger sequencing of genomically integrated STK19 mutant cDNAs to confirm the presence of the following amino acid substitutions in STK19: ΔRNAPII (D154A-T158A-V161A-N162A-V168A-W174A-W175A), ΔUVSSA (K203A-Y204A), and ΔCSA (R72A-T73A-D76A-D98A). **b.** Nuclear localization of TY1-tagged STK19 WT or point-mutants of STK19 designated ΔRNAPII, ΔCSA, and ΔUVSSA expressed in XPC/*STK19-*dKO cells. Scale bar, 20 μm. **c.** Western blot analysis of chromatin-bound TY1-tagged STK19 WT or mutant STK19 proteins ΔRNAPII, ΔCSA, and ΔUVSSA expressed in XPC/*STK19-*dKO cells. TFIIH subunits XPB and p62 serve as a loading control. **d.** Quantification of γH2AX levels normalized to trabectedin-treated parental cells within each experiment. All cells are depicted as individual semi-transparent data points with the bar representing the mean of all data points. The individual means of three biological replicates are depicted as solid circles with black lines. **e.** Endogenous RNAPII-S2 immunoprecipitation (IP) in XPC/*STK19-*dKO RPE1 cells stably expressing STK19 WT or mutant STK19 proteins ΔRNAPII, ΔCSA, and ΔUVSSA. The association of TY1-tagged STK19 proteins with the TCR complex was monitored after UV irradiation (12 J/m^2^, 1 h recovery). TCR complex proteins were analyzed with the indicated antibodies. The data shown represent at least three independent experiments. **f.** Representative images of RNAPII-S2 signal in (GGR-proficient) RPE1 cells with the indicated genotype at 0 h or 1 h after local UV damage (100 J/m^2^) marked by CPD staining. Cells were treated with DRB immediately after UV irradiation to prevent new RNAPII molecules from moving into gene bodies. Scale bar, 10 μm. **g.** Quantification of **f**. The RNAPII-S2 mean pixel intensity at sites of local damage was divided by the mean pixel intensity outside local damage in the same cell nucleus to correct for a decrease of RNAPII-S2 signal due to a general RNAPII run-off caused by DRB. A ratio below 1 signifies that RNAPII-S2 loss in the local damage is faster than the general run-off outside the local damage. All cells are depicted as individual semi-transparent data points with the bar representing the mean of all data points. The individual means of three biological replicates are depicted as solid circles with black lines.

**Figure S7.**
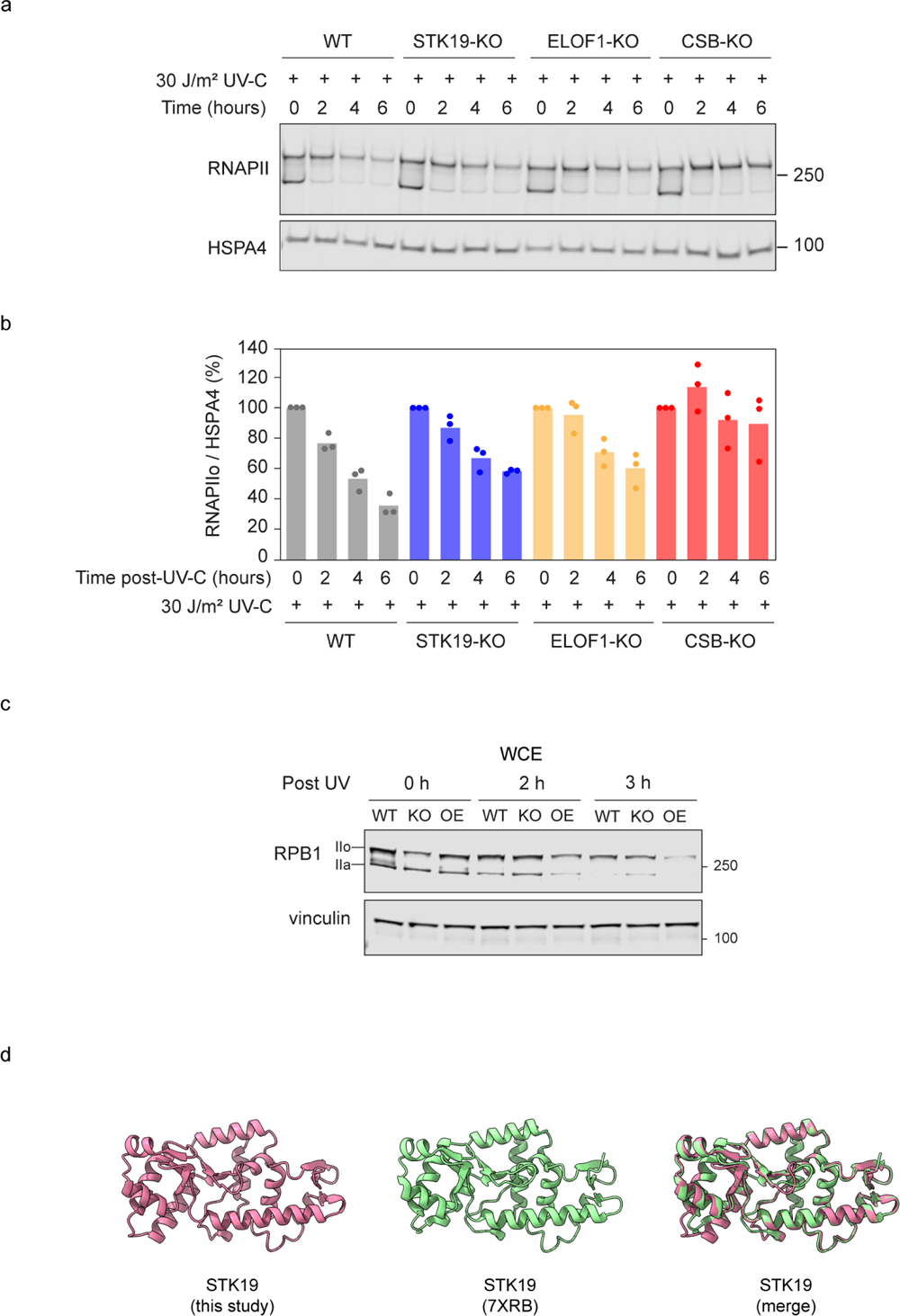
STK19-deficient cells show delayed RPB1 degradation. **a.** Western blot analysis of RPB1 protein levels in RPE1 cells with the indicated genotype that were untreated or irradiated with UV-C (30 J/m^2^, 2, 4, 6 h recovery). HSPA4 serves as a loading control. **b.** Quantification of RPB1 levels normalized to the HSPA4 loading control from three independent experiments. Bars represent the mean. Each circle represents the mean of an independent experiment. **c.** Western blot of a time course after UV irradiation showing the amount of total unphosphorylated (IIa) and phospholylated (IIo) RPB1 in WT, STK19-KO and overexpressing (OE) STK19 Flp-In T-REx HEK293. Vinculin is used as loading control. **d.** Comparison between the STK19 cryo-EM structure determined here compared to the STK19 crystal structure determine by Li et al. (7XRB) (Li et al., 2024).

**Supplementary Table 1:** Cell lines.

| Cell lines | Source |
| --- | --- |
| FlpIn TRex HEK293 (WT) | Thermo Fisher Scientific Cat#R78007 |
| FlpIn TRex HEK293 *CSB*-KO | (Tufegdzcic Vidakovic et al., 2020) |
| FlpIn TRex HEK293 STK19 over-expression (OE) | (Rodriguez-Martinez et al., 2020) |
| FlpIn TRex HEK293 *STK19*-KO | (Rodriguez-Martinez et al., 2020) |
| FlpIn TRex HeLa (WT) | Kind gift from Stephen Taylor |
| FlpIn TRex HeLa *STK19*-KO | (Rodriguez-Martinez et al., 2020) |
| MRC5VA (WT) | Svejstrup lab |
| RPE1-iCas9 (WT) | (van der Weegen et al., 2021) |
| RPE1-iCas9 *CSB*-KO (1-15) | (van der Weegen et al., 2021) |
| RPE1-iCas9 *CSB*-KO (1-15) + *ELOF1*-KO (2-20) | This study |
| RPE1-iCas9 *CSB*-KO (1-15) + *HMCES*-KO (16) | This study |
| RPE1-iCas9 *CSB*-KO (1-15) + *HMCES*-KO (18) | This study |
| RPE1-iCas9 *CSB*-KO (1-15) + *XRCC1*-KO (5) | This study |
| RPE1-iCas9 *CSB*-KO (1-15) + *XRCC1*-KO (6) | This study |
| RPE1-iCas9 *ELOF1*-KO (2-16) | (van der Weegen et al., 2021) |
| RPE1-iCas9 *HMCES*-KO (7) | This study |
| RPE1-iCas9 *HMCES*-KO (9) | This study |
| RPE1-iCas9 *STK19*-KO (7) | This study |
| RPE1-iCas9 *STK19*-KO (7) + STK19-TY(WT) | This study |
| RPE1-iCas9 *STK19*-KO (7) + STK19-TY(WT) + *CSA*-KO (12) | This study |
| RPE1-iCas9 *STK19*-KO (7) + STK19-TY(WT) + *CSA*-KO (3) | This study |
| RPE1-iCas9 *STK19*-KO (7) + STK19-TY(WT) + GFP-UVSSA | This study |
| RPE1-iCas9 *XPC*-KO (5) | (van den Heuvel et al., 2023) |
| RPE1-iCas9 *XPC*-KO (5) + Flag-Bio-DDB2 | This study |
| RPE1-iCas9 *XPC*-KO (5) + Flag-Bio-DDB2 + *CSA*-KO (24) | This study |
| RPE1-iCas9 *XPC*-KO (5) + Flag-Bio-DDB2 + *CSA*-KO (27) | This study |
| RPE1-iCas9 *XPC*-KO (5) + *STK19*-KO (8) | This study |
| RPE1-iCas9 *XPC*-KO (5) + *STK19*-KO (8) + Flag-Bio-DDB2 | This study |
| RPE1-iCas9 *XPC*-KO (5) + *STK19*-KO (8) + STK19-TY(WT) | This study |
| RPE1-iCas9 *XPC*-KO (5) + *STK19*-KO (9) | This study |
| RPE1-iCas9 *XPC*-KO (5) + *STK19*-KO (9) + Flag-Bio-DDB2 | This study |
| RPE1-iCas9 *XPC*-KO (5) + *STK19*-KO (9) + STK19-TY(D154A-T158A-V161A-N162A-V168A-W174A-W175A) (ΔRNAPII) | This study |
| RPE1-iCas9 *XPC*-KO (5) + *STK19*-KO (9) + STK19-TY(K203A-Y204A) (ΔUVSSA) | This study |
| RPE1-iCas9 *XPC*-KO (5) + *STK19*-KO (9) + STK19-TY(R72A-T73A-D76A-D98A) (ΔCSA) | This study |
| RPE1-iCas9 *XPC*-KO (5) + *STK19*-KO (9) + STK19-TY(WT) | This study |
| RPE1-iCas9 *XRCC1*-KO (7) | This study |
| RPE1-iCas9 *XRCC1*-KO (9) | This study |

**Supplementary Table 2:**
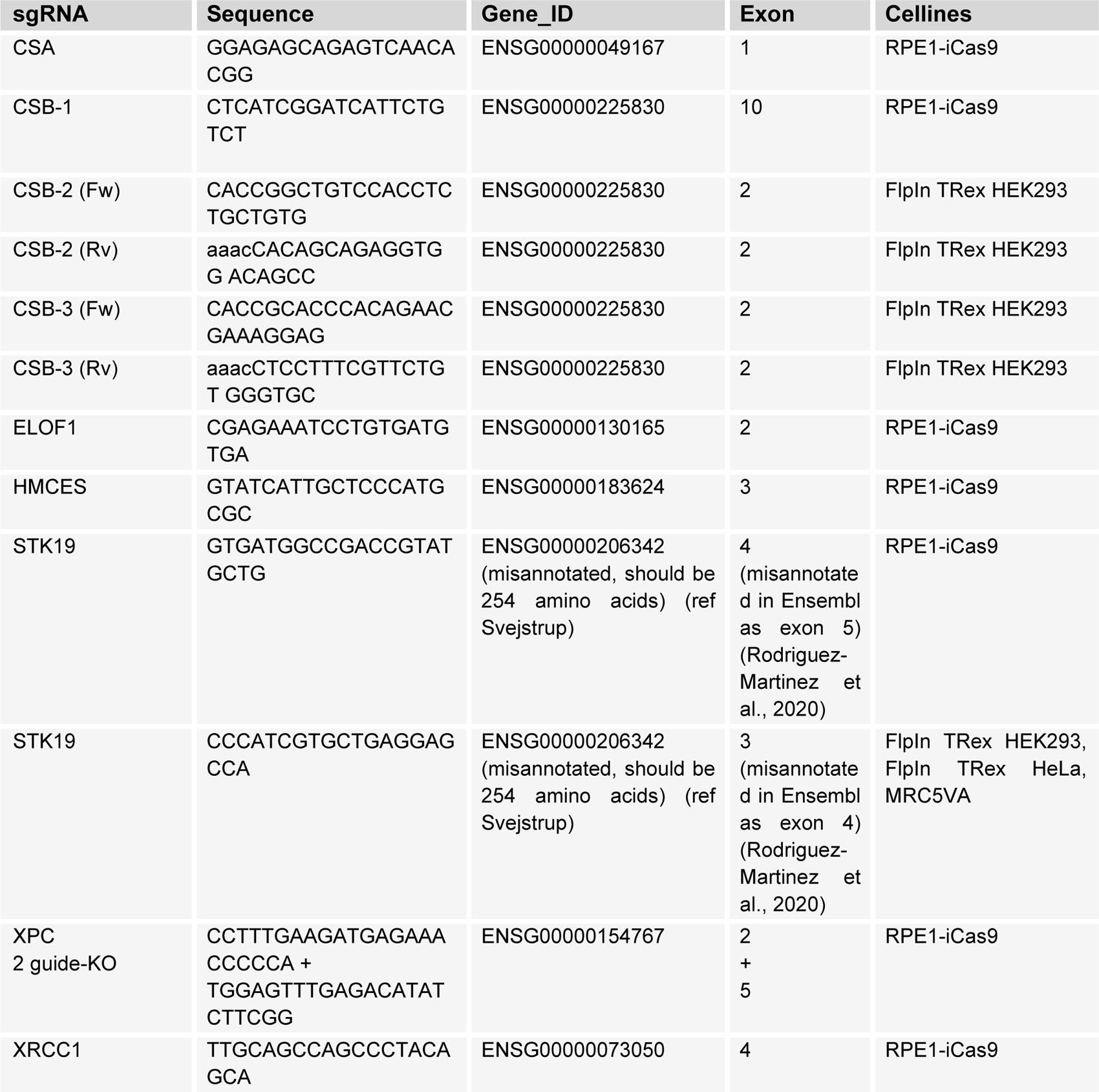
sgRNAs.

**Supplementary Table 3:** Plasmids.

| Plasmid | Origin | Identifier |
| --- | --- | --- |
| pFRT/TO | Gift from Markus Landthaler | (Gregersen et al., 2019) |
| pFRT/TO STK19-Flag | (Rodriguez-Martinez et al., 2020) |  |
| pFRT/TO STK19-Flag-GFP | (Rodriguez-Martinez et al., 2020) |  |
| PGK-3xFlag-Bio-DDB2-IRES-Puro | This study | pML#352 |
| PGK-EGFP-C1-IRES-puro | (Kokic et al., 2024) | pML#244 |
| PGK-EGFP-N1-IRES-puro | (Kokic et al., 2024) | pML#245 |
| PGK-GFP-UVSSA-IRES-puro | (Kokic et al., 2024) | pML#277 |
| PGK-STK19(154A-158A-161A-162A-168A-174A-175A)-TY-IRES-Puro | This study | pML#436 |
| PGK-STK19(203A-204A)-TY-IRES-Puro | This study | pML#437 |
| PGK-STK19(72A-73A-76A-98A)-TY-IRES-puro | This study | pML#419 |
| PGK-STK19-TY-IRES-puro | This study | pML#267 |
| PKG-TY-IRES-puro | This study | pML#260 |
| pOG44 | Thermo Fischer Scientific | Cat#V600520 |
| pU6-gXPC:PGK-puro-2A-tagBFP | Mission library Sigma | sgML#013/014 |
| pX458 | Addgene #48138 | pML#006 |
| pX458-(Cas9-2A-GFP)-sgCSA-3 | (Kokic et al., 2024) | pML#219 |
| pX458-(Cas9-2A-GFP)-sgELOF1-2 | (Kokic et al., 2024) | pML#186 |
| pX458-(Cas9-2A-GFP)-sgHMCES | This study | pML#234 |
| pX458-(Cas9-2A-GFP)-sgSTK19 | This study | pML#214 |
| pX458-(Cas9-2A-GFP)-sgXRCC1 | This study | pML#383 |

**Supplementary Table 4:**
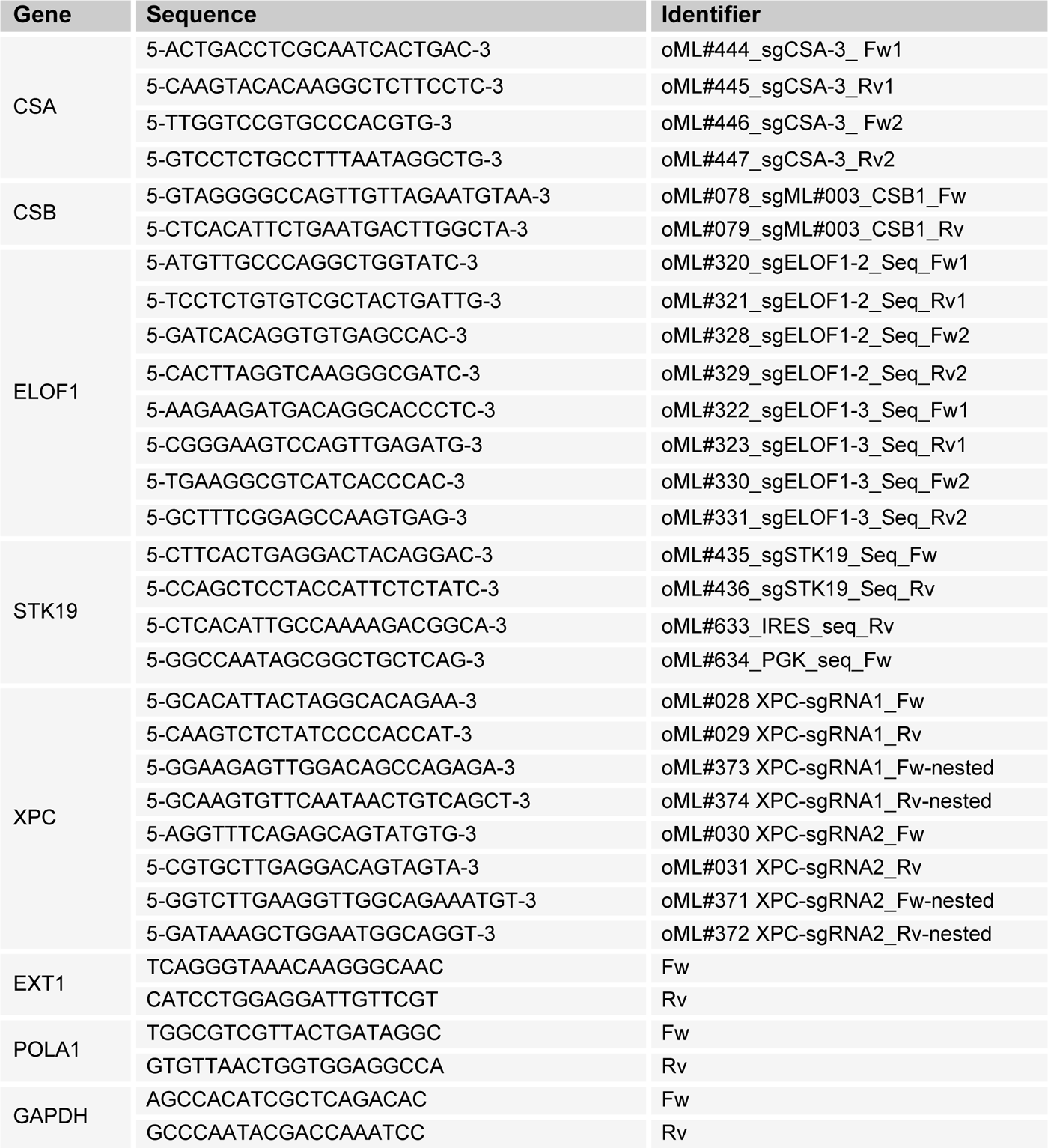
Sequencing and qPCR primers.

**Supplementary Table 5:**
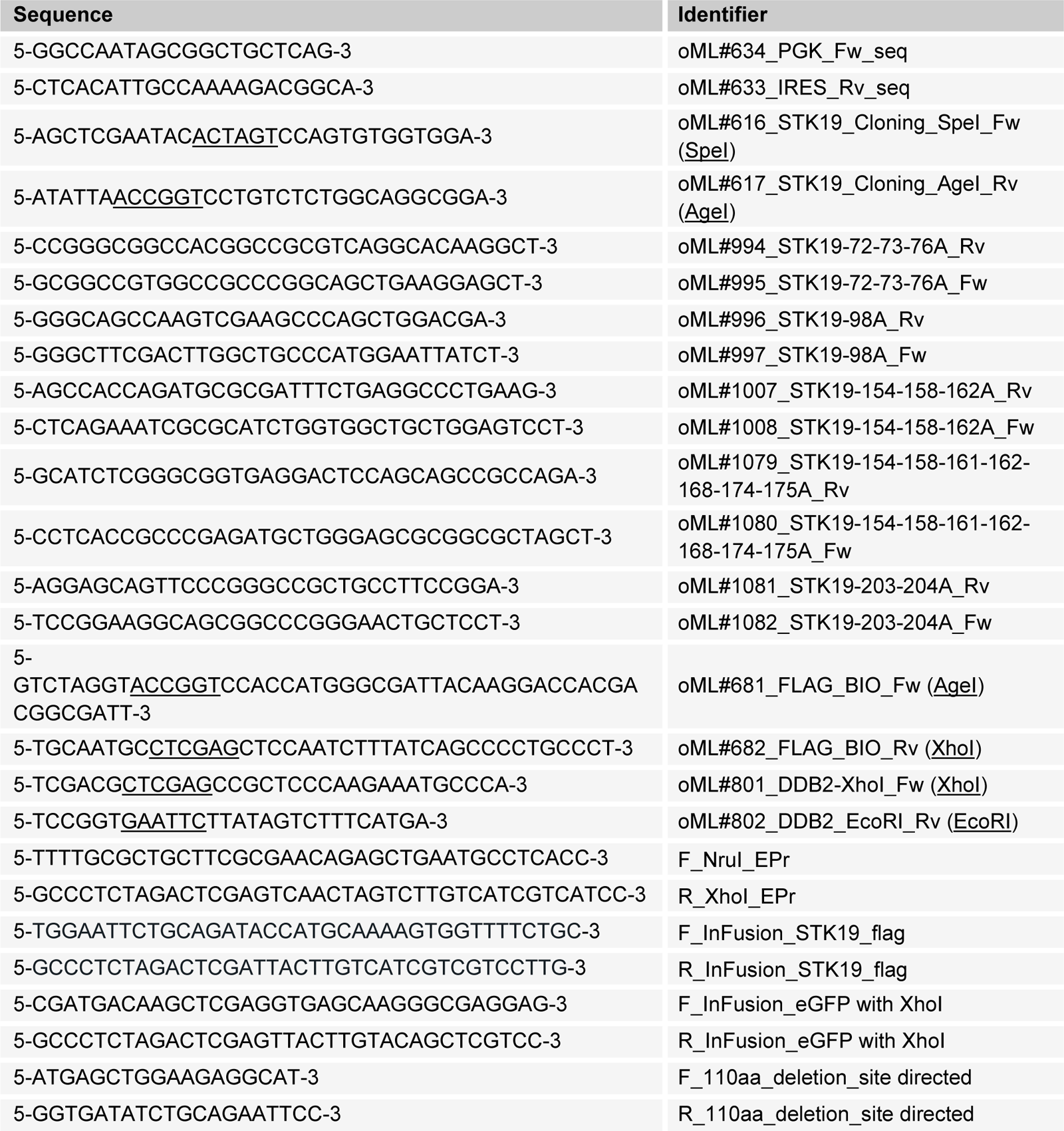
Primers for cloning.

**Supplementary Table 6:** Antibodies.

| Antibody | Host | Company (reference) | Use | Identifier |
| --- | --- | --- | --- | --- |
| Anti-bio | Recombinant protein | Thermo, S11223 | IF: 1:1000 | aML#193 |
| ATF3 | Rabbit | Abcam | WB: 1:1000<br>IF: 1:100 | aML#064 |
| α-Tubulin | Mouse | Sigma, #T6199 (DM1A) | WB: 1:1000 | aML#008 |
| CPD | Mouse | Cosmo Bio, CAC-NM-DND-001 | IF: 1:500 | aML#020 |
| CSA/ERCC8 | Mouse | Santa Cruz, #sc-376981 (D2) | WB: 1:500 | aML#025 |
| CSA/ERCC8 | Rabbit | Abcam, #137033 (EPR9237) | WB: 1:500 | aML#028 |
| CSB/ERCC6 | Rabbit | Santa Cruz, #sc-25370 (H-300) | WB: 1:300 | aML#003 |
| CSB/ERCC6 | Rabbit | Bethyl Laboratories, #A301-345A | WB: 1:600 | aML#187 |
| Flag | Mouse | Sigma-Aldrich (Cat#F3165) |  |  |
| GFP | Mouse | Roche, #11814460001 (7.1 and 13.1) | WB: 1:1000 | aML#011 |
| GFP | Rabbit | Abcam, #ab290 | WB: 1:1000 | aML#044 |
| Histone H3 | Rabbit | Abcam (Cat#ab1791) |  |  |
| HMCES | Rabbit | Atlas Antibodies, HPA044968 | WB:1:500 | aML#214 |
| HRP | Mouse | Santa Cruz |  |  |
| HRP | Rabbit | Jackson ImmunoResearch |  |  |
| HSPA4 | Rabbit | Novus Biologicals, NBP1-81696 | WB: 1:1000 | aML#114 |
| Mouse Alexa 555 | Goat | Thermo fisher Scientific, A-21424 | IF: 1:1000 | aML#015 |
| Mouse Alexa 647 | Goat | Thermo fisher Scientific, A-21235 | IF: 1:1000 | aML#017 |
| Mouse IgG (H+L) CF770 | Goat | Biotium, VWR #20077 | WB: 1:10000 | aML#009 |
| p62/GTF2H1 | Mouse | Santa Cruz, #sc-48431 (G10) | WB: 1:500 | aML#099 |
| p89/XPB/ERCC3 | Mouse | Millipore, #MABE1123 | WB: 1:2000 | aML#101 |
| phospho-H2A.X Ser139 | Mouse | Merck, #05-636 (JBW301) | IF: 1:1000 | aML#161 |
| Rabbit IgG (H+L) CF680 | Goat | Biotium, VWR #20067 | WB: 1:10000 | aML#010 |
| RNAPII (CTD) | Mouse | Cell signalling (4H8; #2629) |  |  |
| RNAPII N-terminus | Rabbit | Cell Signaling, mAb #14958 (D8L4Y) | WB: 1:1000 | aML#252 |
| RNAPII-S2 | Rabbit | Abcam, #ab5095 | WB: 1:1000 | aML#024 |
| RNAPII-S2 | Rat | Millipore, 04-1571 (3E10) | ChIP: 3ug | aML#120 |
| STK19 | Rabbit | Novus Biologicals (Cat#NBP2-33955) |  |  |
| TY | Mouse | Diagenode, C15200054 | WB: 1:1000;<br>IF: 1:6000 | aML#154 |
| Ubiquitin (FK2) | Mouse | ENZO Life Sciences, BML-PW8810-0500 | WB: 1:1000 | aML#102 |
| Ubiquitin (P4D1) | Mouse | Cell Signaling | WB: 1:1000 | aML#192 |
| Vinculin | Mouse | Sigma-Aldrich (Cat#V9131) |  |  |
| XPC | Rabbit | Novus Biologicals, NB100-58801 | WB: 1:2000 | aML#077 |
| XRCC1 | Rabbit | Novus Biologicals, NBP1-87154 | WB: 1:1000 | aML#226 |

**Supplementary Table 7:** Model validation using Molprobity.

| Structure | RNAPII-TCR-ELOF1-STK19 |
| --- | --- |
| PDB / EMD | PDB: 9ER2<br>EMD-19909 |
| Magnification | 120 000 |
| Voltage (kV) | 200 |
| Electron exposure (e / Å<sup>2</sup>) | 39.6 |
| Defocus range (µm) | 0.5-2.5 |
| Pixel size (Å) | 1.23 |
| Symmetry imposed | C1 |
| Initial particle images (no.) | 778 361 |
| Final particle images (no.) | 113 958 |
| Map resolution (Å) | 3.3 |
| FSC threshold | 0.143 |
| Map resolution range (Å) | 2.5-6 |
| Refinement |  |
| Initial model used (PDB code) | 8B3F, AlphaFold |
| Model resolution (Å) | 3.3 |
| FSC threshold | 0.5 |
| Model composition |  |
| Non-hydrogen atoms | 100 220 |
| Protein residues | 6 093 |
| Nucleotide | 106 |
| Ligands | 11 Zn, 2 Mg, BeF<sub>3</sub>, ADP |
| B factors (Å<sup>2</sup>) |  |
| Protein | 95.08 |
| Nucleotide | 163.40 |
| Ligand | 128.37 |
| R.m.s. deviations |  |
| Bond lengths (Å) | 0.012 |
| Bond angles (°) | 0.961 |
| Validation |  |
| MolProbity score | 1.81 |
| Clashscore | 6.46 |
| Poor rotamers (%) | 0.06 |
| Ramachandran plot |  |
| Favored (%) | 92.95 |
| Allowed (%) | 7.05 |
| Disallowed (%) | 0 |

